# Scanning DIA on the ZenoTOF 8600 system records a precursor dimension for every fragment at tenfold higher ion flux

**DOI:** 10.64898/2026.03.12.711261

**Authors:** Tim Heymann, Denys Oliinyk, Lukas Henneberg, Michael B. Lorenz, Nils Eikmeier, Marvin Thielert, Marc Oeller, Louisa Grauvogel, Yaoting Sun, Cole S. Sitron, Bill Loyd, Yves Le Blanc, Nic Bloomfield, Mircea Guna, Adam Carew, Pavel Ryumin, Rob Nieckarz, Ashish Pradhan, Ihor Batruch, Jason Causon, Anjali Chelur, Gordana Ivosev, Katherine Tran, Tatjana Talamantes, Bradley Schneider, James Wighton, Jose Castro-Perez, Matthias Mann

## Abstract

Proteomics needs instruments that combine depth with quantitative precision. Here we describe the ZenoTOF 8600 system, a quadrupole time-of-flight instrument that combines triple-quadrupole ion optics with an optical detector, the Zeno trap and electron-activated dissociation. The MS2 ion current increases tenfold over its predecessor, the ZenoTOF 7600+ system. In scanning quadrupole data-independent acquisition (ZT Scan DIA), every fragment records the Q1 profile of its precursor, so wide isolation windows keep their sensitivity while abundant precursors attain the selectivity of a window about ten times narrower. ZT Scan DIA identifies up to 38% more precursors than variable-window DIA and improves quantitative accuracy. The platform quantifies up to 5,200 protein groups in a single HeLa cell, 5,000 from 1 nL of mouse brain tissue, 10,000 in a single run and 5,700 at 500 samples per day, and targeted assays reach 20 amol on column and quantify S129 phosphorylation of α-synuclein.

## Introduction

Mass spectrometry (MS)-based proteomics has become an indispensable tool for biological and biomedical research, enabling the identification and quantification of thousands of proteins from complex samples^1^. Over the past two decades, advances in instrumentation, chromatography, and computational methods have progressively expanded what is analytically achievable, from early studies identifying hundreds of proteins to current workflows routinely quantifying over ten thousand protein groups from single samples^2,3^. This progress has opened new frontiers, from characterizing the proteomes of individual cells to profiling post-translational modifications at scale, while simultaneously creating demand for instruments that can deliver both the depth required for discovery and the quantitative precision needed for clinical translation and large-cohort studies^4,5^.

Historically, mass spectrometry platforms have presented researchers with a fundamental trade-off. Triple-quadrupole instruments, long the gold standard for targeted quantification, offer exceptional sensitivity and reproducibility but are limited to predetermined analyte panels and lack the accurate mass information needed for confident identification of unknowns. Quadrupole time-of-flight (QTOF) systems provide mass accuracy and spectral quality essential for discovery proteomics but have traditionally lagged behind triple quadrupoles in raw sensitivity. SCIEX introduced the TripleTOF 5600 system in 2010, which combined triple-quadrupole front-end technology with high-resolution TOF analysis, followed by the TripleTOF 6600 system and the development of SWATH data-independent acquisition in 2012^6^. The ZenoTOF 7600 system, launched in 2021, introduced the Zeno trap – a linear ion trap that accumulates and synchronizes the ion arrival time with the pulser to push ions into the TOF analyzer and achieve greater than 90% duty cycle – and electron-activated dissociation for orthogonal fragmentation capabilities^7^.

The ZenoTOF 8600 system represents the next step in this evolution, integrating ion source and transmission technologies from the SCIEX 7500+ system with the versatility of accurate mass analysis with the Zeno trap. Key front-end components from the SCIEX 7500+ system were transferred including the OptiFlow Pro ion source and the DJet and QJet ion guides, while developing a new optical detection system and adapting Mass Guard technology for wide mass range TOF operation. This is combined with the Zeno trap and electron activated dissociation (EAD) cell carried forward from the ZenoTOF 7600 system while advancing scanning DIA (ZT Scan DIA) from the ZenoTOF 7600+ system. Both front-end developments that the ZenoTOF 8600 system inherits have been characterized in isolation: a redesigned ion guide markedly improves ion transfer from atmosphere into the first vacuum stage, and matching the sampling interface to the electrospray plume raises sampling efficiency at microliter and higher flow rates^8,9^. At nanoflow rates, fixed optimal values for key source parameters remove much of the tuning that conventional nanoflow sources require^10^. Together this architecture aims to deliver up to ten-fold greater sensitivity than the previous system while maintaining the resolution, mass accuracy, and scan speed that define modern QTOF performance^11^.

Here we describe the design principles and proteomics performance of the ZenoTOF 8600 system. We compare it with its predecessor, show how scanning quadrupole DIA uses the Q1 dimension to resolve co-isolated precursors that stepped-window DIA cannot, and benchmark the platform for quantitative accuracy in a three-species standard, for single cells and microdissected tissue, at high throughput, and for targeted quantification of peptides and of a disease-relevant phosphosite. Together, these data establish the current capabilities of the platform and illustrate how the combination of triple-quadrupole-derived ion optics with accurate mass versatility can serve applications ranging from basic discovery to clinical research. Prototypes of the ZenoTOF 8600 system were evaluated at the Max Planck Institute of Biochemistry during its development, with results fed back into the design, and all benchmarks in the following sections were acquired and analyzed there.

## Results

### Instrument architecture and sensitivity improvements

The ZenoTOF 8600 system combines established technologies from the SCIEX triple-quadrupole and ZenoTOF platforms with newly developed components optimized for high ion flux operation (Fig. 1a). At the front end, the OptiFlow Pro ion source feeds ions through a sampling orifice enlarged from 0.61 mm to 1.5 mm diameter, increasing the orifice area approximately six-fold compared to the ZenoTOF 7600+ system. Ions enter the DJet ion guide, a tapered dodecapole geometry that efficiently captures ions in the high-pressure region behind the orifice and focuses them into the subsequent QJet ion guide. The QJet ion guide operates in dual-frequency mode, allowing RF frequency to be tuned for optimal transmission of either low or high m/z ions depending on the experiment.

**Figure 1.**
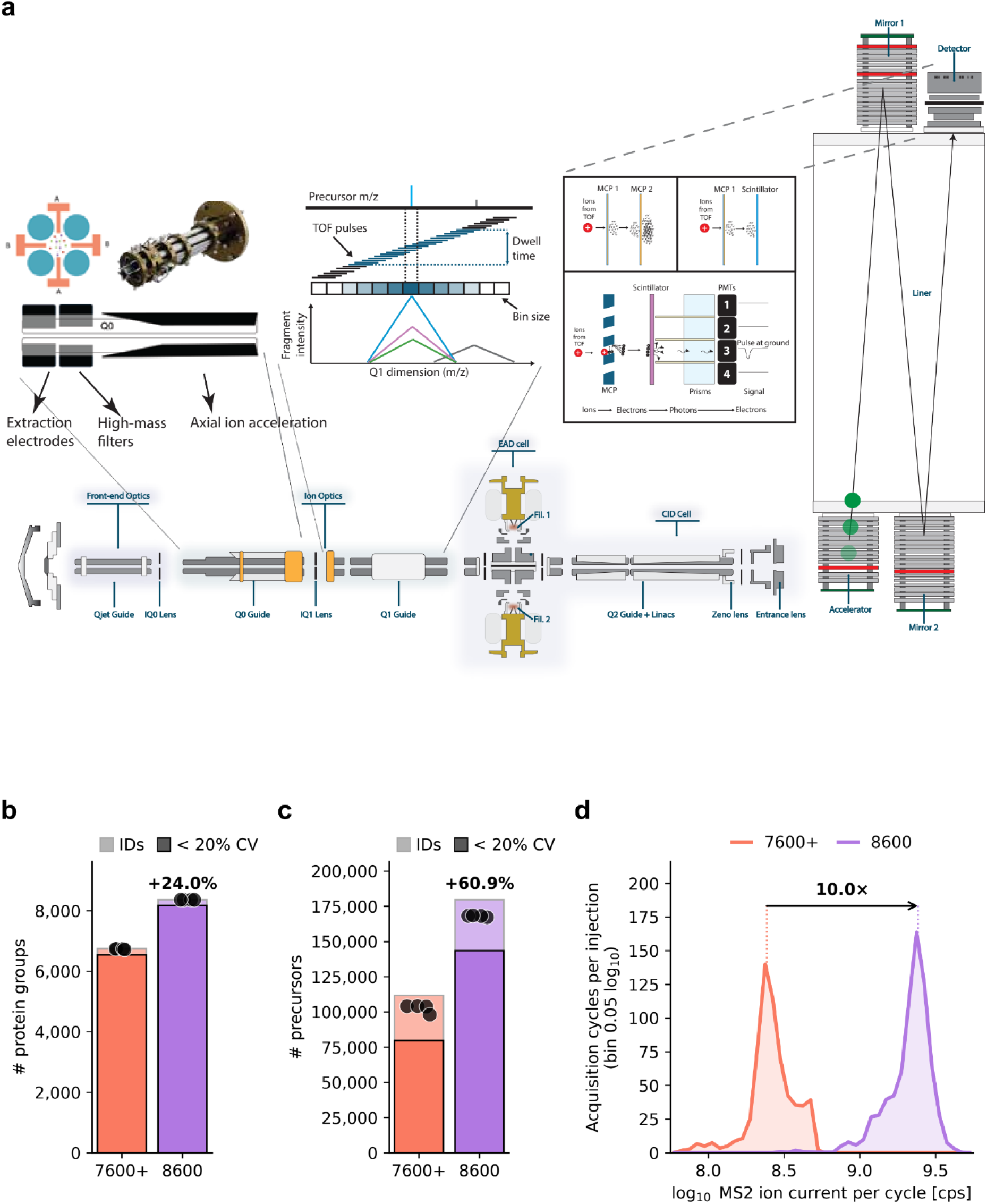
Instrument architecture and sensitivity of the ZenoTOF 8600 system. **a** Schematic of the ZenoTOF 8600 system highlighting the enlarged front-end optics with Mass Guard T-bar electrodes, ZT Scan DIA and the optical (scintillator–photomultiplier) detection system. **b** Protein groups identified from 250 ng HEK293F digest on the ZenoTOF 7600+ and ZenoTOF 8600 systems with the same 60 SPD gradient and Zeno SWATH DIA method (100 variable windows, 12 ms accumulation; n = 4 injections per instrument). Light bars, union of identifications across replicates; dark bars, subset quantified with a coefficient of variation (CV) < 20%; points, individual injections. **c** Same as b for precursors. **d** Distribution of the MS2 ion current per acquisition cycle from the raw files (counts per second) for all cycles within each instrument’s elution window (central 99% of identified precursors), counted per injection (n = 4); dotted lines, modes (median over injections of each injection’s mode); arrow, their ratio.

To maintain performance under sustained high ion current, two protective mechanisms were developed. First, Mass Guard technology employs T-bar electrodes within the Q0 region that apply a DC potential to selectively deflect high-mass contaminants away from the ion beam, functioning analogously to a guard column in liquid chromatography. This filtering protects the Q1 mass analyzer and downstream optics. Second, the traditional dual microchannel plate (MCP) detector was replaced with a hybrid design in which a single MCP feeds a scintillator coupled to four independent photomultiplier tubes. Distributing the ion load across four independent channels extends the linear dynamic range and the lifetime of the detector compared with a single-channel design.

The ZenoTOF 8600 system retains the Zeno trap and electron-activated dissociation (EAD) cell from the ZenoTOF 7600 system. The Zeno trap accumulates ions at the exit of the collision cell and releases them heaviest first, so that ions of all m/z reach the pulser together; this raises the duty cycle to greater than 90%, against the 5–25% typical of conventional orthogonal-injection QTOF designs, with a gain that depends on m/z^12,13^. EAD provides tunable electron-based fragmentation complementary to collision-induced dissociation, enabling enhanced characterization of post-translational modifications and lipid structures^14,15^.

To measure the combined effect of these changes, we analyzed 250 ng of one HEK293F digest on the ZenoTOF 7600+ and ZenoTOF 8600 systems with the same Evosep Eno LC system, column, 60 samples per day (SPD) gradient and Zeno SWATH DIA method (n = 4 each), so that only the mass spectrometer differed. All discovery data in this study were processed with a library-free search in PEAKS Studio 13.5 (Methods)^16^. The ZenoTOF 8600 system identified 24% more protein groups and 61% more precursors than its predecessor (8,368 and 179,779; Fig. 1b, c). Quantification improved as well: 98% of the protein groups had a coefficient of variation (CV) below 20%, with a median protein-group CV of 2.9% against 4.8% on the ZenoTOF 7600+ system (Supplementary Fig. 1a). The precursor intensity distribution shifted upwards by more than half an order of magnitude (Supplementary Fig. 1b, f). Both systems report intensity in counts per second of detected ions, so the increase in MS2 ion current at 600 nL/min, tenfold from peak to peak and 8.8-fold in median (Fig. 1d), is directly comparable and matches the manufacturer’s instrument specification of up to tenfold more transmitted ions^11^. This effectively translated into a median 7.1-fold gain in precursor intensity for the precursors quantified in every injection on both instruments (Supplementary Fig. 1c). The higher signal lowered CVs across the entire abundance range and increased data completeness (Supplementary Fig. 1d, e). Mass accuracy and resolution were maintained at this signal level: across all identified precursors of one injection, the calibrated mass error had a median absolute value of 4.7 ppm for the monoisotopic MS1 peaks and 6.1 ppm for the detected MS2 fragments on the ZenoTOF 8600 system, against 5.3 and 5.8 ppm on the ZenoTOF 7600+ system; the median MS1 resolution was about 38,000 at m/z 780–880 on both, with similar values for MS2 fragments (Supplementary Fig. 1g, h; Methods).

### Scanning quadrupole DIA resolves co-isolated precursors

A key acquisition capability of the ZenoTOF 8600 system is scanning quadrupole DIA (ZT Scan DIA), the commercial implementation of scanning SWATH, which sweeps the quadrupole continuously across the mass range rather than stepping between discrete isolation windows^17–20^. This removes the settling time between windows and encodes the precursor m/z of every fragment in the profile it traces across the Q1 dimension by binning the TOF extractions at a sufficient sampling rate (Fig. 2a). The PEAKS search engine is optimized for ZT Scan DIA data and reads their Q1 dimension through a SCIEX ZT Scan API. To compare the two acquisition modes, we acquired 250 ng HEK293F digest on an Evosep Whisper Zoom 40 SPD gradient with Zeno SWATH DIA and with ZT Scan DIA at different Q1 isolation widths from 1 to 7.2 Th, at a constant 1.48 s cycle matching the SWATH DIA method (n = 4 each). Up to 4.3 Th, every ZT Scan DIA setting identified more precursors than variable-window DIA, with 250,090 at 1 Th against 180,850 in Zeno SWATH DIA, 38% more (Fig. 2b, c). Precursors and protein groups quantified with CV < 20% peaked at 2.8 Th (167,485 and 8,972, against 146,008 and 8,526), which we therefore used as the default.

**Figure 2.**
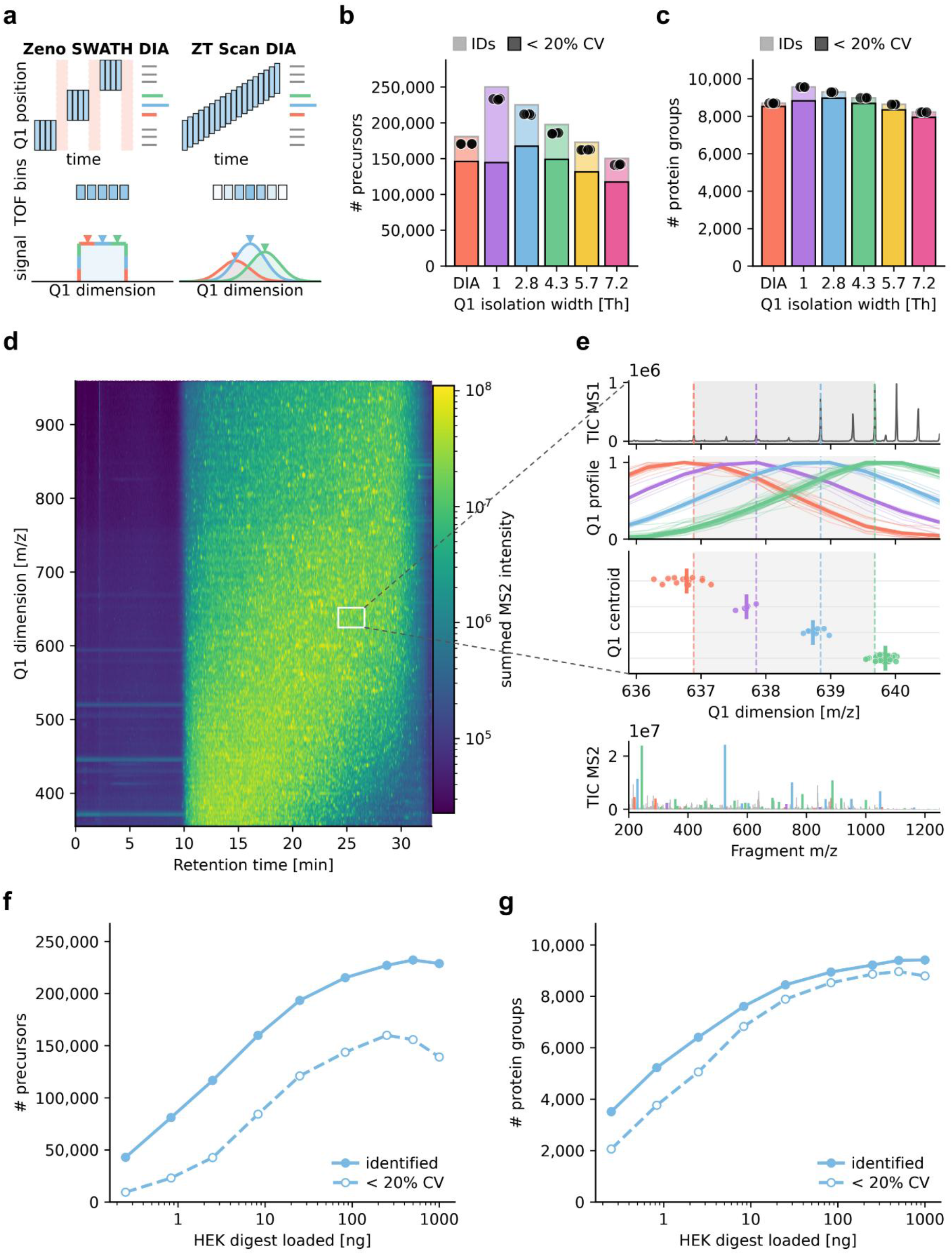
Scanning quadrupole DIA resolves precursors in the Q1 dimension. **a** Principle of Zeno SWATH DIA (left; shaded, quadrupole settling time) and ZT Scan DIA (right), in which every fragment traces a Q1 profile whose position reports its precursor m/z. **b** Precursors identified from 250 ng HEK293F digest (Whisper Zoom 40 SPD) with Zeno SWATH DIA (100 variable windows, 12 ms) and ZT Scan DIA at Q1 isolation widths of 1–7.2 Th (1.48 s cycle); n = 4 injections per method. Light bars, union of identifications across replicates; dark bars, subset quantified with a coefficient of variation (CV) < 20%; points, individual injections. **c** As in b, for protein groups. **d** Summed MS2 intensity across retention time and the Q1 dimension for one 2.8 Th ZT Scan DIA run; every cell is one MS2 spectrum. The white box marks the region of interest shown in e. **e** A 2.8 Th Q1 range (25.4 min, m/z 636.9–639.7) containing four co-eluting precursors. From top: MS1 spectrum; normalized Q1 profiles of fragments unique to each precursor (thin, individual fragments; thick, mean); Q1 centroids of the individual fragments (points) and their median (bar); the chimeric MS2 spectrum of the window with fragments colored by precursor. Grey shading, isolation window; dashed lines, precursor m/z from MS1. **f** Precursors identified (solid) and quantified with CV < 20% (dashed) across a dilution series of HEK293F digest (0.25–1,000 ng) with the 2.8 Th method; n = 4 injections per load. **g** As in f, for protein groups.

Because each fragment inherits the Q1 profile of its precursor, those that are co-isolated in one window can be separated by the centroid of that profile (Fig. 2d, e). We illustrate this with a single 2.8 Th window, in which four co-eluting precursors identified by the search contribute to one chimeric MS2 spectrum. The Q1 profiles of the fragments of each precursor correlate strongly and center on its respective precursor mass. Even the two closest precursors in this example window (0.84 Th apart) are separated by about six times the locating error of the Q1 centroid (σIQR 0.14 Th; Supplementary Fig. 2a). Across all precursors identified by PEAKS, the Q1 centroid often located precursors to a tenth of the isolation window or better, provided that their signal was high enough to correlate the Q1 profiles of their fragments (Supplementary Fig. 2b). Depending on the isolation width, 14–32% of the precursors at or above the median abundance (7,795 to 22,612) had Q1 profiles clean enough to pass a simple criterion (Methods). After calibration of the Q1 axis, which held on peptides not used to fit it and carried over between injections (σIQR 0.140 Th with an axis transferred from another injection, as with the injection’s own; Methods; Supplementary Fig. 2a, c), the Q1 centroid located these precursors with a precision (σ_IQR_) of 0.12 Th at 1 Th and 0.14 Th at 2.8 Th isolation width. This is 3.1-fold (1 Th) to 10-fold (5.7 Th) better than the isolation window alone. Consequently, for precursors located with this precision, the average number of co-isolated precursors fell from 4.8 within a 2.8 Th window to 0.6 within ±1σ of the Q1 centroid (Supplementary Fig. 2d): ZT Scan DIA transmits a wide window yet, when precursors are told apart by the Q1 centroids of their fragments, attains the selectivity of a window about ten times narrower, although each fragment spectrum itself remains chimeric. Because this holds at every window width, a wider isolation window costs far less selectivity than its width suggests: even at 7.2 Th, the average number of co-isolated precursors fell from 12.6 within the window to 1.3 within ±1σ of the Q1 centroid (Supplementary Fig. 2d). For a peptide of interest, the fragments that share its Q1 profile give a precursor-specific, DDA-like spectrum from the DIA data (Fig. 2e). In these analyses, the Q1 dimension locates precursors whose identities are known from the search; how much it contributes to the identifications was not assessed, and deconvolving chimeric spectra without this prior knowledge remains to be implemented.

Across more than three orders of magnitude of input, from 250 pg to 1,000 ng of HEK293F digest, the 2.8 Th method identified 3,500 to 9,400 protein groups, and identifications quantified with CV < 20% peaked at 250 ng for precursors and 500 ng for protein groups (Fig. 2f, g). With a 1 Th method, halving the throughput to 20 SPD with a longer cycle added only 4.6% protein groups, reaching 10,000 (Supplementary Fig. 2e), so the 40 SPD method already captures 96% of the protein groups identified at 20 SPD, and we used 40 SPD in the following experiments.

### Quantitative accuracy in a three-species benchmark

Quantitative accuracy is commonly tested with mixed-species benchmarks, in which proteomes of several organisms are combined at known ratios^21^. We prepared mixtures of commercially available human (K562), yeast and E. coli digests in which the human fraction was kept constant at 50% while yeast and E. coli varied inversely from 45% to 5% across six conditions (Fig. 3a), and analyzed 250 ng of each with ZT Scan DIA (2.8 Th) and with Zeno SWATH DIA on a Whisper Zoom 40 SPD gradient (n = 4). With ‘match between runs’ across the six mixtures, similar numbers of protein groups were identified for each species in every mixture (Fig. 3b). ZT Scan DIA identified 13,290 protein groups and 230,475 peptides across all 24 injections, 10% and 24% more than Zeno SWATH DIA (12,097 and 185,296; Fig. 3c, d).

**Figure 3.**
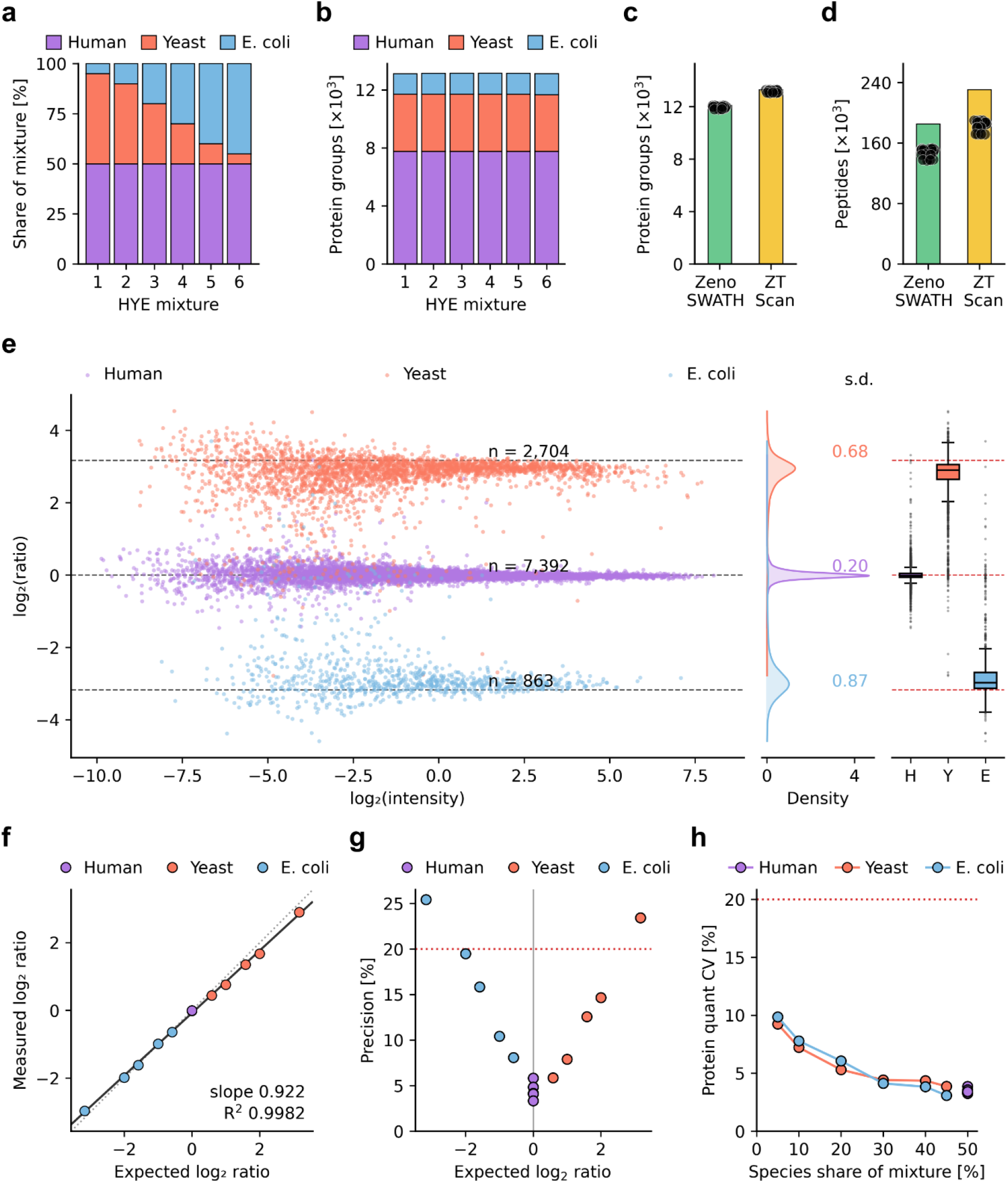
Quantitative accuracy in a three-species benchmark. **a** Composition of the six mixtures (250 ng total per injection): human digest held at 50%, yeast and E. coli varied inversely from 45% to 5%. **b** Protein groups identified per species in each mixture (ZT Scan DIA, union of n = 4 injections, match between runs across mixtures). **c** Protein groups identified with Zeno SWATH DIA (100 variable windows, 12 ms) and ZT Scan DIA across all 24 injections per method; bars, union; points, individual injections. **d** As in **c**, for peptides. **e** log_2_ ratio against log_2_ intensity of protein groups for mixture 1 against mixture 6, colored by species; dashed lines, expected ratios. Right, density of each species’ ratios with their standard deviation, and box plots (center, median; box, interquartile range; whiskers, 1.5 × interquartile range) against the expected ratios (red). **f** Measured against expected log_2_ ratio for five mixture pairs and three species (median over proteins); solid line, linear fit; dotted line, identity. **g** Spread of the ratios between proteins of one species, expressed as (2^σ^ − 1) × 100%, with σ the σ_IQR_ of the log_2_ ratios, for each pair and species. **h** Median protein CV across n = 4 injections against the share of the species in the mixture. Red dotted lines in **g**, **h**, 20%. **e–h**, ZT Scan DIA (2.8 Th); quantification as described in Methods. The Zeno SWATH DIA comparison is shown in Supplementary Fig. 3.

Quantitative accuracy was assessed by comparing measured protein ratios against expected values. For each pair of mixtures, protein intensities were calculated with directLFQ from the precursors quantified in all four replicates of both, without further filtering^22^. Even for the largest designed fold change, nine-fold between mixtures 1 and 6, the three species formed separate ratio distributions centered close to their expected values (Fig. 3e). With Zeno SWATH DIA, the yeast and E. coli distributions lay closer to zero, with median log_2_ ratios of 2.63 and −2.77 against 2.90 and −2.97 for ZT Scan DIA (expected ±3.17; Supplementary Fig. 3a). Across five mixture pairs and three species, measured ratios followed the expected ratios with a slope of 0.92 and R² = 0.998 (Fig. 3f), against 0.84 for Zeno SWATH DIA (Supplementary Fig. 3b). The remaining compression was confined to yeast, whose median ratios reached 75–92% of the expected log_2_ ratios, whereas those of E. coli lay within 10% of expectation in every pair.

Precision was assessed in two ways. Because all proteins of one species share the same designed ratio, the spread between them directly measures quantification error. It was 3–6% for human proteins, whose ratio does not change, and rose with fold change for yeast and E. coli, from 6–8% at 1.5-fold to 23–25% at nine-fold. Over all pairs and species, the median spread was 8.1% for ZT Scan DIA against 9.9% for Zeno SWATH DIA, and lower for ZT Scan DIA in all 15 comparisons (Fig. 3g, Supplementary Fig. 3c). The median replicate CV of individual proteins was 4.0% against 4.6% (lower in 15 of 18 mixture and species comparisons) and depended on the amount of a proteome in the mixture rather than on its species, with yeast and E. coli following one curve (Fig. 3h, Supplementary Fig. 3d).

### Proteomics at low input: single cells and tissue

Single-cell proteomics represents perhaps the most demanding test of instrument sensitivity, as protein content per cell typically falls in the sub-nanogram range^4^. At low input, wider Q1 windows helped moderately compared to the 2.8 Th default (Fig. 2g): on 0.25 ng HEK293F digest, a single-cell equivalent, ZT Scan DIA with a 7.2 Th window identified the most protein groups, slightly more than Zeno SWATH DIA (Supplementary Fig. 4a, b). The enhanced sensitivity mode, a detector setting with increased detector voltage (Methods), further increased precursor intensities 1.76-fold, which improved precursor identifications by 10%, and precursors quantified with CV < 20% by 22% (Supplementary Fig. 4c–e). With both settings, we analyzed individual HeLa cells isolated by image-based sorting with the cellenONE and prepared in an automated oil-based 384-well plate workflow^23^ (Fig. 4a). Single cells yielded a median of 4,466 protein groups, with up to approximately 5,200, while the 20-cell pools approached 5,600, close to the 5 ng bulk digest of approximately the same protein content (Fig. 4b). Processed blanks yielded more than an order of magnitude fewer precursors than single cells (median 1,986 against 30,406), which remained similar when searched without the 20-cell pools and 5 ng digest as references (1,960 against 23,365; Supplementary Fig. 4f, g), indicating a low background of about 2,000 precursors. Excluding all precursors found in the processed blanks reduced the median number of protein groups per single cell by 1.5%. The data of the 20 single cells captured cell-to-cell variation: the cells could be assigned to G1, S and G2/M phases (5, 9 and 6 cells) using protein-based cell-cycle scores (Fig. 4c). Protein abundances spanned more than three orders of magnitude between the 1st and 99th percentiles, with transcription factors such as STAT3, RELA and SP1 quantified up to 3.7 orders of magnitude below housekeeping proteins such as ACTB (Fig. 4d).

**Figure 4.**
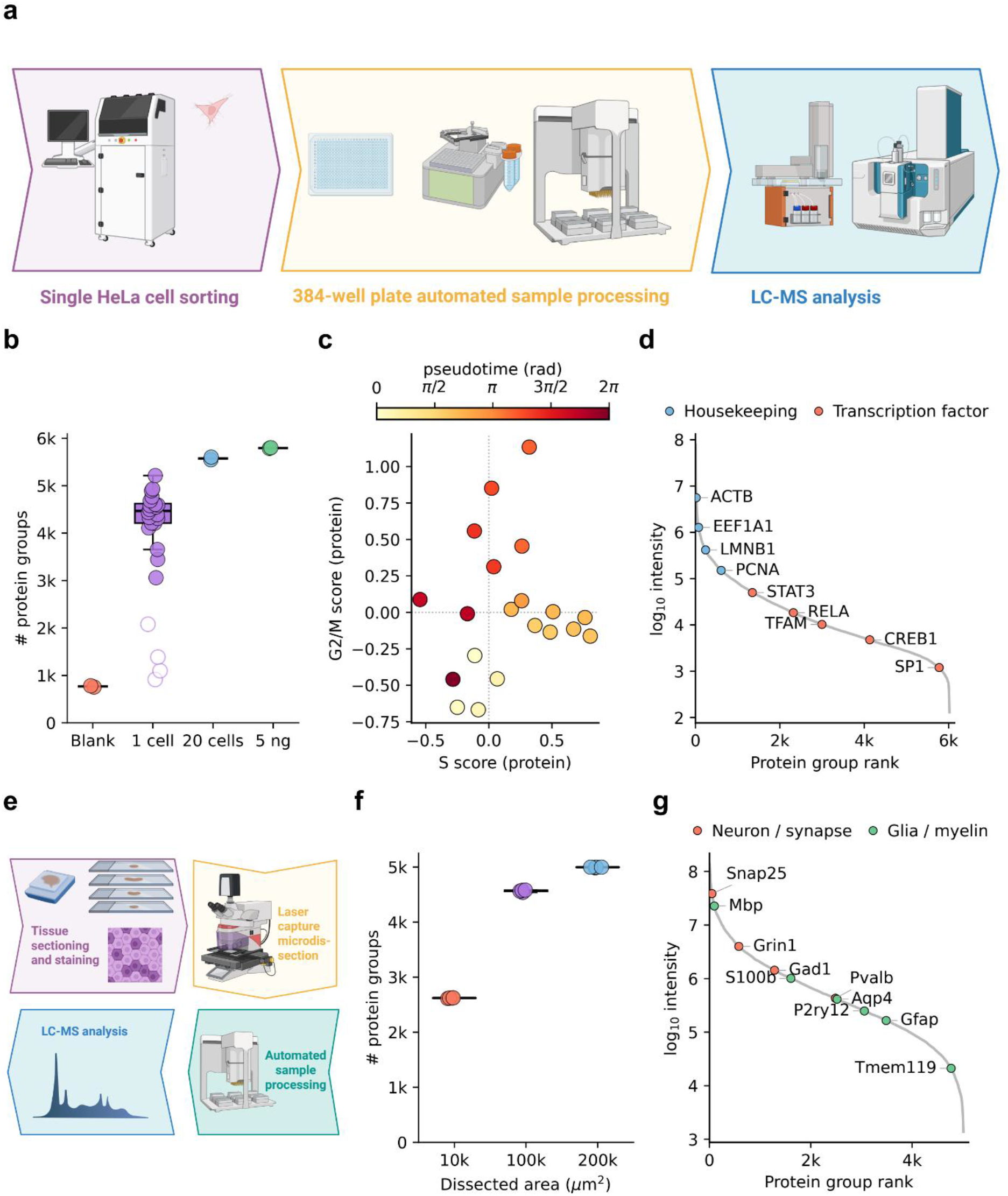
Single-cell and spatial tissue proteomics. **a** Workflow for single HeLa cells: cell sorting, automated sample processing in 384-well plates and LC-MS analysis. **b** Protein groups per run for processed blanks (n = 2), single HeLa cells (n = 20), 20-cell pools (n = 2) and 5 ng HeLa digest (n = 2), acquired with ZT Scan DIA (7.2 Th, Whisper Zoom 40 SPD). Box, median and interquartile range of the single cells; whiskers, 1.5 × interquartile range; lines, median of the groups of two. Four of 24 cells with outlying low precursor counts were excluded (open points; Methods). **c** Cell-cycle scores of the 20 single cells computed from S-phase and G2/M marker proteins, colored by pseudotime angle. **d** Rank–abundance curve of protein groups across the single cells (median log_10_ intensity over the cells in which each protein was quantified), with housekeeping proteins and transcription factors highlighted. **e** Deep Visual Proteomics workflow: tissue sectioning, staining and segmentation, laser microdissection, automated sample processing and LC-MS analysis. **f** Protein groups identified from 10,000, 100,000 and 200,000 µm² of microdissected mouse brain tissue with the method in **b** (n = 4 dissections per area; lines, median). **g** Rank–abundance curve for the 200,000 µm² samples with neuronal and synaptic (orange) and glial and myelin (green) markers highlighted.

Besides single cell proteomics, spatial proteomics of tissue is similarly demanding, because microdissected regions contain little material and cannot be pooled without losing spatial information. Using Deep Visual Proteomics (Fig. 4e), we analyzed laser-microdissected mouse brain sections of 10,000 to 200,000 µm² with the single-cell method^24^. A 10,000 µm² region of a 5 µm section (50 pL) yielded 2,621 protein groups, and 200,000 µm² (1 nL) 5,000 (Fig. 4f), with the proteome spanning five orders of magnitude and markers of neurons, astrocytes, oligodendrocytes and microglia covering more than three, from Snap25 to the microglial marker Tmem119 (Fig. 4g).

### High-throughput proteomics

Fast, selective acquisition on short gradients has recently been shown to give deep proteome coverage at high throughput^2^. However, on gradients shorter than 10 minutes, a stepped quadrupole has little time to cover the whole precursor range, whereas a continuously scanning quadrupole loses no time to settling between windows, which makes scanning DIA particularly advantageous on short gradients. We acquired 250 ng human, yeast and E. coli digests at 200, 300 and 500 SPD (6.4, 3.95 and 2.3 min gradients) and, at each throughput, ran Zeno SWATH DIA and ZT Scan DIA at the same 0.6 s cycle so that the acquisition scheme was the only difference within a pair (n = 4; Fig. 5a, b). The cycle time was chosen to maximize quantifiable precursors while sampling peaks of approximately one second width about twice across their full width at half maximum (FWHM) (Supplementary Fig. 5a–d), a sampling density close to that of the three-species benchmark (about 2.5 points across the FWHM), in which we had recovered ratios accurately (Fig. 3).

**Figure 5.**
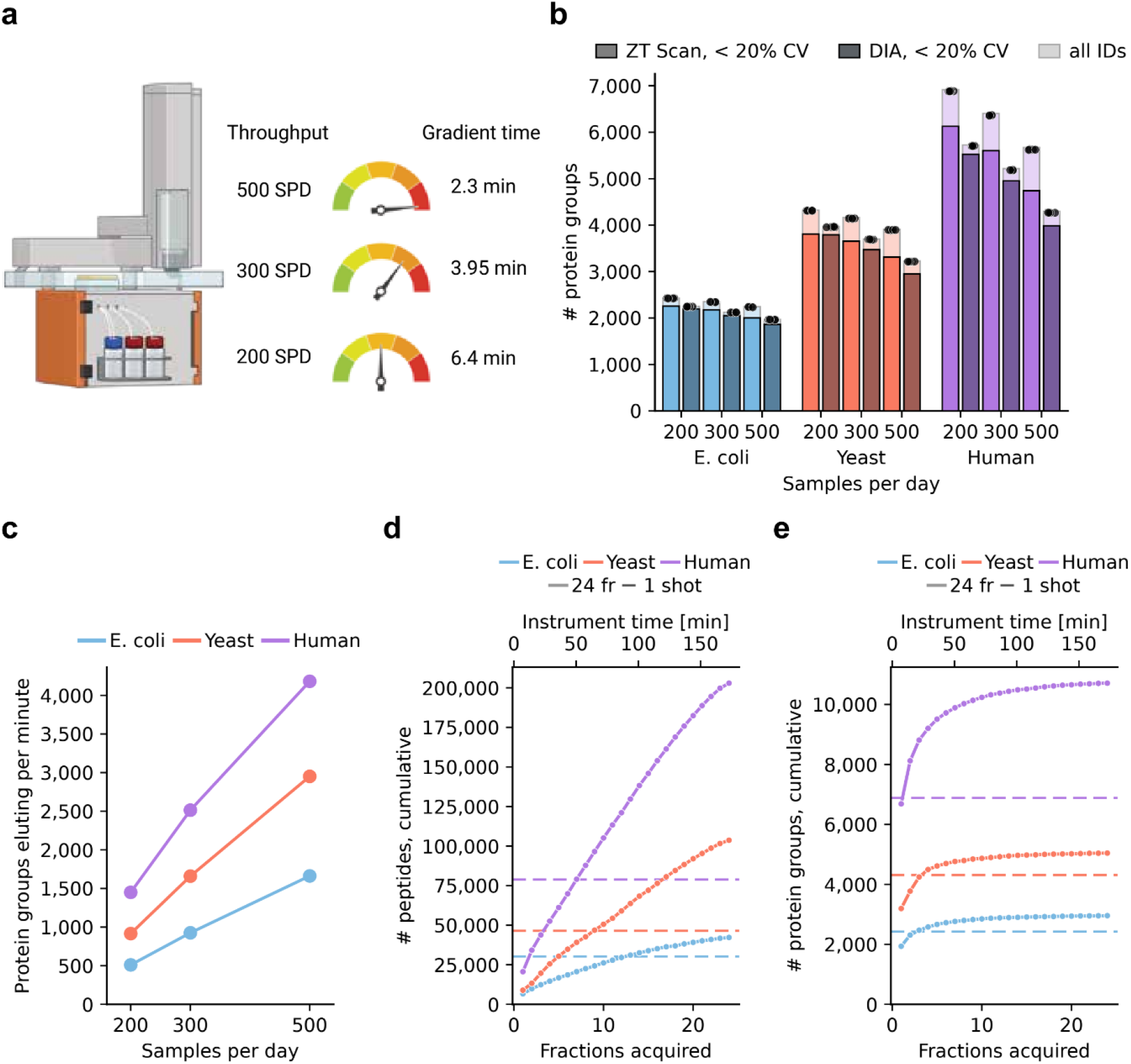
Proteome depth at high throughput. **a** Evosep gradients used for high throughput data acquisition: 500, 300 and 200 samples per day (SPD; 2.3, 3.95 and 6.4 min gradients). **b** Protein groups identified from 250 ng of HEK293F, yeast and E. coli digests at each throughput with ZT Scan DIA (2.1 Th, 0.6 s cycle) and Zeno SWATH DIA (80 variable windows, 5 ms, 0.6 s cycle); n = 4 injections per condition; in each pair, ZT Scan DIA brighter and Zeno SWATH DIA darker. Light bars, union of identifications across replicates; dark bars, subset quantified with a coefficient of variation (CV) < 20%; points, individual injections. **c** Protein groups identified per minute of elution window with ZT Scan DIA, the window being the retention-time span of the central 99% of identified precursors in each run. **d** Cumulative peptides identified across 24 high-pH reversed-phase fractions of 100 µg HEK293F, yeast and E. coli digest acquired at 200 SPD with ZT Scan DIA (top axis, instrument time); dashed lines, single-shot injections at 200 SPD from **b** (separate acquisitions). **e** As in **d**, for protein groups.

As expected, the advantage of ZT Scan DIA grew as the gradient shortened (Fig. 5b). For human digest, it identified 6,900, 6,400 and 5,700 protein groups at 200, 300 and 500 SPD, 21% to 32% more than Zeno SWATH DIA (5,700, 5,200 and 4,300), and it also identified more protein groups for yeast and E. coli at every throughput. At 500 SPD, this amounts to more than 4,000 human protein groups per minute of elution (Fig. 5c).

For deeper coverage, we combined ZT Scan DIA with offline fractionation. We separated 100 µg each of HEK293F, yeast and E. coli digest into 48 high-pH reversed-phase fractions, concatenated them to 24 and acquired each at 200 SPD at a load of ∼250 ng per injection. In less than three hours of instrument time, this yielded 10,700 human protein groups and 203,000 human peptides, about 1.6- and 2.6-fold more than single-shot analysis at 200 SPD, for 24 times the injected peptide amount (6 µg against 250 ng) and 24 times the instrument time (Fig. 5d, e). Protein identifications approached saturation within a few fractions while peptide identifications continued to rise and sets of 6 or 12 fractions reached a given depth with fewer injections while covering a shallower depth in total (Fig. 5d, e; Supplementary Fig. 5e).

### Targeted quantification and phosphosite analysis

Biomarker validation, pharmacokinetic studies and clinical assays require targeted measurement of specific analytes with high sensitivity and quantitative accuracy. In MRM^HR^ (high-resolution multiple reaction monitoring), the quadrupole isolates each precursor within a window of about 1 Th (unit resolution) and the TOF records all of its fragments at high resolution, so a targeted assay keeps the full MS/MS spectrum for confirming the analyte.

We evaluated targeted quantification with the Promega 6×5 LC-MS/MS Peptide Reference Mix spiked into 250 ng HEK293F digest, in which five isotopologues of each peptide span four orders of magnitude, from 2 amol to 20 fmol, in one injection (Methods). With scheduled MRM^HR^ at 200 SPD at unit resolution, the five peptides retained on Evotips yielded well-resolved chromatographic peaks, while the remaining peptide was too polar to be retained on Evotips (Fig. 6a). Calibration curves were linear across the quantifiable range (weighted R² = 0.964–0.997 across the five peptides; LGFTDLFSK in Fig. 6b). Three of the five peptides were quantified at 20 amol and all five at 200 amol, and the median limits of detection and quantification, extrapolated from the calibration curves, were 5.5 and 16.6 amol on column. Back-calculated amounts deviated by less than 20% from nominal for 69 of 72 quantified points, with a median recovery of 99.4% (Fig. 6c), and the median CV across injections decreased from 8.8% at 20 amol to 1.6% at 20 fmol (Fig. 6d).

**Figure 6.**
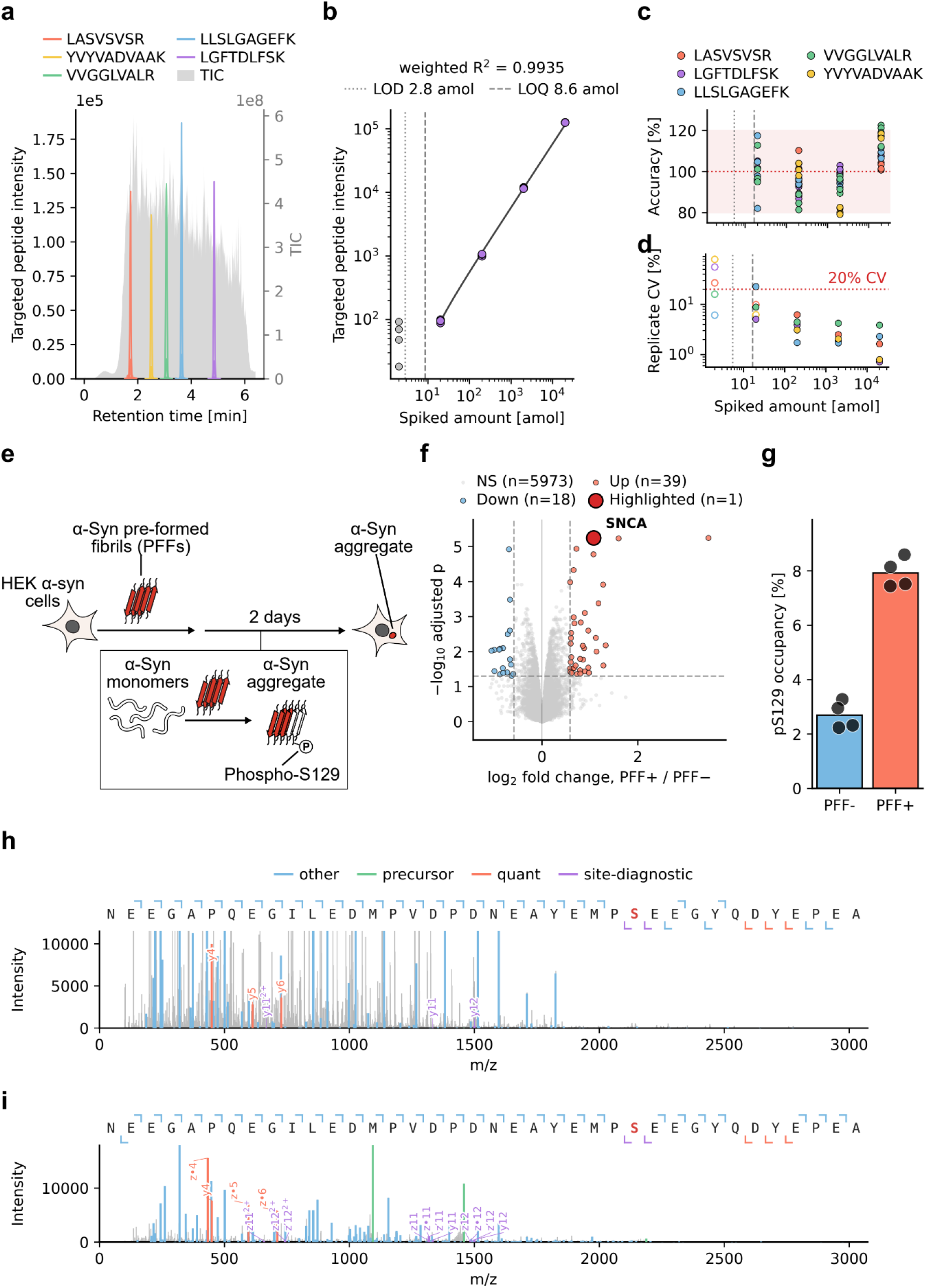
Targeted quantification and phosphosite analysis by MRM^HR^. **a** Extracted ion chromatograms of the five peptides of the Promega 6×5 standard, each present as five isotopologues from 2 amol to 20 fmol, spiked into 250 ng HEK293F digest and acquired in one MRM^HR^ injection at 200 SPD; grey, total ion current (right axis). **b** Calibration curve of LGFTDLFSK (n = 4 injections); line, 1/x²-weighted linear fit; grey points, level below the fitted range; dotted and dashed lines, limit of detection (LOD) and quantification (LOQ). **c** Accuracy of the back-calculated amounts for all quantified levels of the five peptides; shaded band, 80–120%; red dotted line, 100%. **d** CV across n = 4 injections; open symbols, levels not accepted (Methods); red dotted line, 20%. Dotted and dashed lines in **c**, **d**, median LOD and LOQ of the five peptides. **e** HEK293T cells expressing α-synuclein treated with pre-formed fibrils (PFF) for two days to induce aggregation and S129 phosphorylation. **f** Protein-level fold changes between PFF-treated (PFF+) and untreated (PFF−) cells (ZT Scan DIA, 2.1 Th, 0.6 s cycle, 200 SPD; n = 4 injections of one digest per condition). Moderated two-sided t-test with Benjamini–Hochberg correction; dashed lines, |log_2_ fold change| = 0.585 and adjusted P = 0.05. **g** pS129 occupancy of α-synuclein determined by MRM^HR^ (EAD) with heavy-isotope-labeled standards; bars, mean; points, repeated injections of one digest per condition (n = 4). h, i CID (h) and EAD (i) MS2 spectra of the heavy-labeled α-synuclein phosphopeptide NEEGAPQEGILEDMPVDPDNEAYEMP(pS)EEGYQDYEPEA (4+), summed across the chromatographic peak; grey, unmatched peaks; z•, radical z ions. Ions are colored by role: quantifier (EAD z4, y4, z5, z6; CID y4–y6), site-diagnostic (y/z11 and y/z12, bracketing S129), precursor, other; y axes are scaled to the assay ions (full scale in Supplementary Fig. 7c, d).

This sensitivity is enabled by efficient ion transmission even at narrow Q1 isolation widths. Of the four resolution settings for MRM^HR^ (high, unit, low and open), narrower ones provide selectivity for closely spaced precursors at the cost of transmission. Measured across the calibrants of the Promega 6×5 standard and comparing the same transitions for the isolated precursor isotope, unit resolution transmitted a median of 59% of the signal obtained with the open setting, high resolution 46%, and low resolution as much as the open setting (Supplementary Fig. 6a–c). Unit resolution therefore offers a practical compromise for most targeted applications and was used throughout. Quantification from the summed MS2 fragment ions is far more sensitive and selective than the precursor isotopes in the MS1 scans of the same injections, which were measurable only down to 2 fmol, one hundred-fold higher (Supplementary Fig. 6d–h). This difference likely reflects that Zeno pulsing is not used for MS1 scans and that the MS1 ion beam is attenuated by dynamic ion transmission control (ITC), which keeps the detector linear at the much larger ion flux of a full-range MS1 scan; MS2 scans sample only a small part of the mass range.

Targeted MRM^HR^ methods allow characterization and validation of challenging analytes requiring maximum sensitivity and selectivity such as post-translational modifications (PTMs) that regulate protein function, localization, and interactions^25^. Modified peptides are often sub-stoichiometric relative to their unmodified counterparts, and confident site localization requires high-quality fragmentation spectra^26^. We applied the capabilities of the ZenoTOF 8600 system to phosphorylation of α-synuclein at serine 129 (pS129), a modification closely linked to the pathological aggregation of α-synuclein in Parkinson’s disease^27,28^. HEK293T cells stably expressing A53T α-synuclein were treated with pre-formed fibrils (PFF) to induce α-synuclein aggregation and the associated S129 phosphorylation^29–31^ (Fig. 6e). Global proteomic analysis by ZT Scan DIA of one PFF-treated and one untreated culture, each injected four times, found 57 proteins that differed beyond technical variation (39 higher, 18 lower), including a 2.1-fold increase of α-synuclein itself (Fig. 6f). Because the cells were treated with recombinant α-synuclein fibrils, part of this increase reflects internalized PFF.

A targeted MRM^HR^ assay with heavy-isotope-labeled standards for the unmodified and phosphorylated C-terminal peptide quantified the stoichiometry of the modification directly. The pS129 occupancy, the phosphorylated fraction of the peptide calculated from the light-to-heavy ratios of both forms (Methods), increased from 2.7% in untreated to 7.9% in PFF-treated cells, a 2.9-fold increase (Fig. 6g), consistent with the established role of this modification in α-synuclein aggregation pathology^27,28^. PFF seeding is known to produce S129-phosphorylated α-synuclein aggregates in cultured cells, so the increase follows the induced aggregation, although the measured occupancy in this case is that of the pooled endogenous and the unphosphorylated internalized protein from the PFF treatment^29,31^. The discovery ZT Scan DIA data of the same samples already yielded a site occupancy and a similar increase (3.7-fold; Supplementary Fig. 7a), which the targeted EAD assay validated with higher selectivity and more informative fragmentation (Fig. 6h, i). The discovery data gave higher absolute occupancies (3.2% and 11.7%, against 2.7% and 7.9% in the targeted assay) since it is less selective than the targeted method and more prone to interference, particularly in the light channel (Supplementary Fig. 7b). Additionally, CID generated fewer of the singly charged fragment ions used for quantification than EAD, even though Zeno pulsing was disabled for EAD. Like most large peptides at higher charge states, this 4+ peptide fragments more productively with electron-based activation than with CID^32^. Still, CID and EAD spectra of the phosphopeptide provided complementary information in this case. CID efficiently produced b- and y-type ion series at high sensitivity for sequence identification (Fig. 6h), whereas EAD produced mainly c- and z-type ions that preserved the labile phosphate and covered the full length of the large C-terminal peptide, confirming the phosphorylation site and peptide identity with high confidence (Fig. 6i, Supplementary Fig. 7c, d). In both spectra, the y11 and y12 ions (and z11 and z12 in EAD) differed by 167.00 Da, the mass of phosphoserine, while y8–y11 were observed unmodified, localizing the phosphate to S129 rather than to the neighboring acceptors Y125, Y133 or Y136. The site-diagnostic ions were also present for the endogenous (light) phosphopeptide in PFF-treated cells: y11 and z11 at their unmodified and y12 and z12 at their phosphorylated masses, at the same light-to-heavy ratio as all other y8–y13 and z8–z13 ions (Supplementary Fig. 7b).

## Discussion

The ZenoTOF 8600 system was built on a simple premise: the ion optics that make triple-quadrupole instruments the most sensitive mass spectrometers should work as well in an accurate mass instrument, provided the detector can keep up with the higher ion current. Our data show that this premise holds. The enlarged front end and the optical detector raised the number of detected ions about tenfold over the predecessor. Together with the scanning quadrupole, this gives the instrument three properties that distinguish it from current accurate mass instruments: very high sensitivity, the capacity to handle very high ion currents, and a recorded precursor dimension for every fragment.

The first two properties, very high sensitivity and the capacity for a large ion current, change what limits an experiment. When ions are plentiful, the constraint shifts from collecting enough signal to using the available signal well. Narrower isolation windows, shorter gradients and, in future, multiplexed samples all spread a fixed ion supply over more measurements, a trade-off that only an instrument accepting more ions can afford. The higher signal lowered CVs across the whole abundance range rather than only adding identifications, and on the shortest gradients, where a stepped quadrupole loses most time to settling, ZT Scan DIA identified about a third more protein groups than Zeno SWATH DIA.

The third property is conceptually important, even though a precursor dimension has been recorded before with scanning SWATH, SONAR, synchro-PASEF and midiaPASEF^17,19,33,34^. A stepped quadrupole collects no information about which precursor produces which fragments in its Q1 isolation window, so all co-isolated precursors share one fragment spectrum. A scanning quadrupole records a Q1 profile for each fragment correlating with its precursor, and the centroid of this profile localizes fragments to precursors to a few tenths of a Thomson, often within less than one tenth of the Q1 isolation window (Supplementary Fig. 2a). The selectivity is therefore set by the precision of the Q1 centroid rather than by the window width, and for sufficiently abundant precursors it is narrower than the 2 Th windows of narrow-window DIA or the 1–2 Th windows typical of DDA methods^2^. This selectivity comes without the overhead of stepped DIA, whose quadrupole settles at every window transition by design: the continuously scanning quadrupole spends its whole cycle acquiring data, which makes ZT Scan DIA an efficient way to generate highly selective data. Unlike ion mobility, which separates by collision cross section and constrains m/z only through its correlation with it, the Q1 dimension reports the precursor m/z directly.

PEAKS, which is optimized for ZT Scan DIA data, uses the Q1 dimension for identification, and its contribution there remains to be quantified. For any peptide of interest, the fragments that share its Q1 profile form a precursor-specific, DDA-like spectrum that can be extracted, and the same principle allows assignment of diagnostic ions such as glycan oxonium ions to their precursors and other fragment ions prior to identification and extends to small molecules^35–38^. Doing this systematically could help search engines score chimeric DIA spectra and, in future, support de novo sequencing and the search for variant and unexpected peptides.

These capabilities do not come at the expense of the established strengths of a QTOF. In a three-species benchmark, ZT Scan DIA recovered expected ratios with lower spread than variable-window DIA. Single cells and nanoliter tissue regions reached depths in the range of current single-cell and spatial proteomics studies^39–41^. Offline fractionation extended human proteome coverage beyond ten thousand protein groups in less than three hours. In targeted mode, MRM^HR^ reached attomole sensitivity with the full fragment spectrum retained, and EAD localized a disease-relevant phosphosite, so the same instrument covers discovery, high-throughput and targeted work, from sample screening to validation.

Several aspects will need further work. The use of the Q1 dimension in data analysis can be greatly expanded in the future. Data processing must keep pace with acquisition, particularly at hundreds of samples per day. The phosphorylation example demonstrates targeted PTM analysis, but phosphoproteomics and other PTM workflows at scale remain to be benchmarked, as does performance on endogenous analytes in clinical specimens, where interference and matrix effects are larger than with synthetic standards. Finally, further improvements of the ion path are possible, which together with the Q1 dimension make the platform suited to proteomics at much larger scale.

## Methods

### Cell culture

HEK293F cells were maintained in suspension culture in FreeStyle 293 Expression Medium in 100–2,000 mL Erlenmeyer flasks with vented caps at 37 °C and 8% CO₂ on an orbital shaker platform at 125 rpm. Cell viability was monitored by trypan blue exclusion and maintained above 90% throughout all experiments. For experiments, cells were cultured to a density of 2.0 × 10⁶ cells/mL, washed twice with cold TBS, pelleted at 200 × g for 10 min, snap-frozen, and stored at −80 °C until use.

HEK293T TetOff-α-syn-A53T cells (RRID: CVCL_D4BW) were cultured at 37 °C and 5% CO₂ in DMEM containing 10% FBS and 1% penicillin-streptomycin supplemented with 2 μg/mL blasticidin S^42^. Cell pellets were collected by trypsinization, washed twice with cold PBS, and frozen at −80 °C before use. HeLa cells (ATCC, S3 subclone) were grown in DMEM with 20 mM glutamine, 10% FBS and 1% penicillin-streptomycin to ∼80% confluency, harvested with 0.25% trypsin-EDTA and processed as HEK293F cells. All cell lines were routinely tested for mycoplasma contamination.

### Escherichia coli

E. coli Rosetta BL21 colonies (LB agar, 37 °C overnight) were pre-cultured in 2 mL LB (10 g/L tryptone, 5 g/L yeast extract, 10 g/L NaCl; 4 h, 37 °C, 180 rpm), diluted 1:100 into 50 mL LB in 250 mL baffled flasks and grown at 37 °C and 220 rpm to mid-exponential phase (OD_600_ ≈ 0.6–0.8). Cells were harvested (4,000 × g, 10 min, 4 °C), washed twice with ice-cold 50 mM Tris-HCl pH 7.5, snap-frozen and stored at −80 °C.

### Saccharomyces cerevisiae

S. cerevisiae strain S288C colonies (YPD agar: 1% yeast extract, 2% peptone, 2% glucose, 2% agar; 2 days, 30 °C) were pre-cultured overnight in 2 mL YPD (30 °C, 180 rpm), and 50 mL YPD in 250 mL baffled flasks was inoculated to a starting OD_600_ of 0.1 and grown at 30 °C and 200 rpm to mid-exponential phase (OD_600_ ≈ 0.8–1.0). Cells were harvested (3,000 × g, 5 min, 4 °C) and processed as above.

For lysis, cell pellets were resuspended in lysis buffer (50 mM TEAB pH 8.5, 40 mM CAA, 10 mM TCEP, 2% SDC) and boiled at 95 °C for 10 min with shaking at 1,500 rpm. Lysates were then subjected to high-energy tip sonication (30 pulses, 20% duty cycle), boiled again at 95 °C for 10 min with shaking at 1,500 rpm, and centrifuged at maximum speed for 5 min to remove insoluble debris. Protein concentration was determined by a tryptophan fluorescence assay.

### αAggregation of α-synuclein in HEK293T cells

α-Synuclein pre-formed fibrils (PFFs) were produced based on an established protocol^43^. Briefly, 5 mg/mL recombinant A53T α-synuclein was centrifuged (100,000 × g, 1 h, 4 °C) and the supernatant agitated at 900 rpm and 37 °C for 24 h. A detailed protocol can be found at https://doi.org/10.17504/protocols.io.btynnpve. For aggregation, HEK293T TetOff-α-syn-A53T cells were seeded at 250 cells/mm². The next day, PFFs were diluted 1:20 in PBS, sonicated in a BioRuptor Plus (25 cycles, 5 s on/5 s off), added to Opti-MEM and Lipofectamine 3000 pre-mixed for 5 min (final 50:3:20, v/v), incubated for 10 min and added dropwise at 1 μL sonicated PFFs per 10 mm² of well; controls received PBS instead. The medium was replaced with DMEM the next day, and cells were harvested two days after treatment as above. A detailed protocol can be found at https://doi.org/10.17504/protocols.io.eq2lyjbbplx9/v1.

### Evotip Pure sample loading

Evotip Pure tips were prepared by washing with 20 μL EvoB (0.1% FA in ACN), priming in 1-propanol for 15 s, and washing with 20 μL EvoA (0.1% FA in water). For loading, 100 μL EvoA was added to the Evotip, into which the digested sample was then pipetted. After loading, tips were washed with 20 μL EvoA and re-wetted with 100 μL EvoA. All centrifugation steps were performed at 700 × g for 1 min.

### Sample preparation

Cell lysates were diluted to 0.5 mg/mL in 50 mM TEAB containing 0.01% DDM and digested with trypsin and LysC (10 ng/μL each) for 2 h at 37 °C with shaking at 1,200 rpm. Digestion was quenched by addition of TFA to a final concentration of 0.5% (v/v), diluting samples to 0.25 mg/mL. Precipitated SDC was removed by centrifugation at maximum speed for 1 min, and the resulting supernatant was loaded onto Evotip Pure tips as described above.

### Sample preparation for single-cell proteomics

Harvested HeLa cells were resuspended in ice-cold PBS to a concentration of 200 cells/µL for an oil-based 384-well workflow^23^. A 384-well plate was pre-loaded with 400 nL lysis buffer per well using the Mantis liquid dispenser (Formulatrix). Lysis buffer contained 100 mM triethylammonium bicarbonate (TEAB), 0.2% n-dodecyl-β-D-maltoside (DDM), 10 ng/µL Lys-C (Wako), and 20 ng/µL Trypsin Platinum (Promega). Single cells were then dispensed into the pre-loaded plate using the cellenONE system (Cellenion, France) at 10 °C and elevated humidity, with sorting parameters set to a cell diameter of 22–26 µm and an elongation factor < 2; 20-cell pools received 20 cells and processed blanks no cell. The plate was incubated at 50 °C for 90 min. After digestion, peptides were acidified by addition of 5 µL of 1% trifluoroacetic acid (TFA) and the plate was cooled to 4 °C. Peptides were loaded onto preconditioned Evotip Pure tips (Evosep Biosystems) using the Bravo liquid handling robot (Agilent), followed by a well wash with 4 µL EvoA, which was also transferred to the Evotip to maximize peptide recovery. Evotips were conditioned as described above, but with two wash steps each of EvoB and EvoA instead of one.

### Mouse tissue preparation

Animals used were bred for scientific purposes at the Max Planck Institute of Biochemistry Animal Facility, and the research in this project does not involve experiments on animals (as defined by law). All animals were sacrificed by CO₂ euthanasia prior to removal of tissue in accordance with the European Commission Recommendations for the euthanasia of experimental animals (Part 1 and Part 2). Breeding, housing, and euthanasia of the animals were fully compliant with all German (the German Animal Welfare Act) and EU (Directive 2010/63/EU) applicable laws and regulations concerning care and use of laboratory animals.

An eight-week-old male C57BL/6J mouse was euthanized and transcardially perfused with rinsing solution (100 mM Tris pH 8.5, 200 mM NaCl, 20 mM EDTA); the brain was removed, washed, and fixed overnight in 4% (w/v) paraformaldehyde. The tissue was then dehydrated with 1 h incubations in 70%, 80%, 90% and 100% ethanol and cleared with two 1 h incubations in xylene before paraffin embedding in disposable molds.

### Deep Visual Proteomics sample preparation

A 5-µm section was cut from a formalin-fixed, paraffin-embedded (FFPE) mouse brain block using a microtome (Thermo Fisher Scientific) and mounted onto a UV-treated 4.0-µm PPS membrane slide (MicroDissect GmbH). The section was deparaffinized and rehydrated (xylene, then 100%, 90%, 75% and 30% ethanol and ddH₂O, twice for 3 min each) and stained with hematoxylin and eosin. Tissue areas of 10,000, 100,000 and 200,000 µm² were collected into a 384-well plate (Eppendorf) using an LMD7 laser microdissection system (Leica Microsystems) with a 20× or 40× objective. Laser cutting was performed in middle-pulse mode (power 55, aperture 1, speed 10, final pulse −1, middle pulse count 2, head current 40%, pulse frequency 1,050, offset 100). Dissected tissue was lysed in 6 µL 50 mM TEAB, 0.01% DDM and 5 mM TCEP at 95 °C for 1 h. Digestion was initiated by adding 2 µL of 4 ng/µL Lys-C and 6 ng/µL trypsin and carried out for 4 h at 37 °C, then quenched with 0.9 µL 10% TFA using a Mantis liquid dispenser (Formulatrix). Peptides were loaded onto Evotips according to the manufacturer’s instructions.

### Commercial digests and standards

Lyophilized commercial tryptic digests of K562 (Promega), yeast (Promega), and E. coli (Waters) were reconstituted to a final concentration of 0.5 mg/mL by sequential addition of 40 μL EvoB (0.1% formic acid in acetonitrile) followed by 160 μL EvoA (0.1% formic acid in water). Digests were further diluted in EvoA containing 0.01% DDM and loaded onto Evotip Pure tips, either individually or combined at the indicated ratios, as described above. The Promega 6×5 LC-MS/MS Peptide Reference Mix was dissolved in 20% acetonitrile with 0.1% formic acid, diluted 1:100 and loaded at 20 fmol (most abundant isotopologue) onto the Evotip together with 250 ng HEK293F digest.

### Liquid chromatography

All samples except those acquired on the ZenoTOF 7600+ system for Fig. 1 were acquired on a ZenoTOF 8600 system (SCIEX) with a pre-commercial, research-grade version of SCIEX OS, coupled to an Evosep Eno LC system (Evosep Biosystems), either in the micro-flow configuration with 500, 300 and 200 SPD on a 4 cm × 150 µm PepSep column and 100 and 60 SPD on an 8 cm × 150 µm PepSep column, or in the nano-flow configuration with the Whisper Zoom 40 and 20 SPD gradients on an Aurora Elite XS column (15 cm × 75 µm, 1.7 µm C18; IonOpticks). For Fig. 1, both systems received the same HEK293F digest batch and amount through the same LC system, column and 60 SPD gradient (600 nL/min). Unless stated otherwise, 250 ng of digest was injected. For the input series, HEK293F digest was injected at 0.25, 0.83, 2.5, 8.3, 25, 83, 250, 500 and 1,000 ng (n = 4 per amount), and each amount was searched separately.

### Mass spectrometry

The ZenoTOF 8600 (research-grade SCIEX OS) and ZenoTOF 7600+ systems (SCIEX) were operated with the column oven at 50 °C. In the micro-flow configuration (500–60 SPD), the 1–10 µL electrode was in the top port and the calibration probe in the front port (gas 1 20 psi, gas 2 60 psi, 300 °C, curtain gas 40 psi, 4,500 V). In the nano-flow configuration (Whisper Zoom gradients), the nano probe with the IonOpticks column was in the front port and the nano calibration probe, connected to the calibrant delivery system, in the top port (heaters 200 °C, gas 2 40 psi, nano gas 20 psi, curtain gas 40 psi, nano cell 300 °C, 2,000 V). Enhanced sensitivity mode, a detector setting with increased detector voltage, was used for all acquisitions on the ZenoTOF 8600 system except those of Fig. 1, the mode-off arm of Supplementary Fig. 4c–e and all MRM^HR^ acquisitions, which were run in the default LDR detector mode; the ZenoTOF 7600+ system has no equivalent setting.

Zeno SWATH DIA methods used 100 variable windows of 4–11 Th between 359 and 961 m/z (Supplementary Table 1), designed with the dynamic binning of pyDIAid from the 246,563 precursors identified by gas-phase fractionation of 250 ng HEK293F digest on the ZenoTOF 8600 system (six 40 SPD injections with 2 Th windows, each covering 200 Th between 300 and 1,500 m/z; Supplementary Fig. 8), 12 ms accumulation time and dynamic collision energy, with a precursor scan from 360 to 1,500 m/z and 50 ms accumulation^44^. For the 200, 300 and 500 SPD comparisons, 80 variable windows of 5–14 Th between 359 and 961 m/z (Supplementary Table 2), designed in the same way, with 5 ms accumulation were used, giving a 0.6 s cycle. In all Zeno SWATH DIA methods, Zeno trap pulsing was activated on demand.

ZT Scan DIA methods covered 360–960 m/z with dynamic collision energy, with Zeno trap pulsing always active. On the Whisper Zoom 40 SPD gradient, Q1 isolation widths of 1, 2.8, 4.3, 5.7 and 7.2 Th were acquired at a 1.48 s cycle time, and 5.7, 7.2, 10.1 and 14.5 Th for the low-input comparison; the 1 Th method was also run on a 20 SPD gradient at a 2.4 s cycle. The 2.8 Th method was used for the input series and the three-species benchmark, and the 7.2 Th method for single cells and microdissected tissue. For 200, 300 and 500 SPD, a 2.1 Th method with a 0.6 s cycle was used; at 500 SPD, methods of 3.3, 2.6, 2.1 and 1.8 Th with cycle times of 0.4, 0.5, 0.6 and 0.7 s were compared.

MRM^HR^ methods used the source settings above, unscheduled or scheduled from a reference run of the same sample and gradient. CID of the 6×5 standard used the default dynamic collision energy for 2+ ions and 50 ms accumulation. For the α-synuclein phosphopeptide, EAD was performed with 10 ms reaction time, 15 eV kinetic energy and 6,500 nA electron beam current with 50 ms accumulation time, and CID in a separate scan as above, without a scan schedule. Zeno trap pulsing was activated on demand for CID and disabled for EAD. The Q1 resolution series was acquired at 100 SPD, with one injection of the 6×5 standard per resolution setting at a 10-fold higher amount (20 amol to 200 fmol per isotopologue).

### Data analysis

All discovery data were searched library-free in PEAKS Studio 13.5 (Bioinformatics Solutions) using the human, mouse, yeast and E. coli FASTA files according to sample content^16^. Peptides of 7–45 amino acids and charge states 1–4 were searched with carbamidomethylation of cysteine as a fixed modification and methionine oxidation and protein N-terminal acetylation as variable modifications (maximum one variable modification, up to two missed cleavages), and results were filtered with the PEAKS default settings, 1% FDR at the precursor level in each run and 1% FDR at the protein-group level, so that only precursors of accepted protein groups were retained. Each condition was searched separately, except that all single cells, each acquisition method of the three-species benchmark and the PFF experiment were each searched as one job. In the three-species benchmark, match between runs was enabled across the six mixtures.

Single HeLa cells were searched library-free, together with the 20-cell HeLa pools (approximately 5 ng) and two injections of 5 ng bulk HeLa digest as references, with match between runs (all shown in Fig. 4b); the single cells and processed blanks were also searched without the references (Supplementary Fig. 4f, g).

### Identification and quantification metrics

Search results were processed from the PEAKS feature-level export with custom Python code (Python 3.12; numpy 2.3, pandas 2.3, scipy 1.16, statsmodels 0.14, alphabase 1.7, alpharaw 0.4.8, directlfq 0.3.3). Precursors were defined as modified sequence and charge, peptides as modified sequence, and protein groups as reported by PEAKS. An entity was identified in a run if its normalized area was non-zero; per condition, the union across injections and the per-injection counts are reported. Protein quantities were the PEAKS protein-group areas unless stated otherwise. CVs were calculated on linear intensities across the replicates of a condition for entities with at least three values, without imputation; entities with CV < 20% form the quantifiable subset. For the instrument comparison, the per-precursor intensity gain was computed for precursors quantified in all four injections on both instruments as the difference of the per-instrument median log_2_ normalized areas. The raw MS2 ion current was compared on the summed intensity per acquisition cycle within each instrument’s elution window, the retention-time span containing the central 99% of the precursors identified in its search (0.5th–99.5th percentile). The peak-to-peak ratio is that of the modes of the logarithmic per-cycle distributions (median over injections of each injection’s mode, from a Gaussian kernel density with Silverman’s robust bandwidth). The median ratio is that of the medians over injections of each injection’s median. Mass accuracy and resolution (Supplementary Fig. 1g, h) were measured in the raw data of one injection per instrument for all identified precursors. Each precursor was located by at least three co-eluting fragments; its MS1 spectra were summed between the half-maximum points of its chromatographic peak and its MS2 spectra across the scans of its isolation window (each sample combined with the closest sample of every other spectrum within 1 ppm, at most one per spectrum), and the m/z of each monoisotopic precursor peak and of each b or y fragment peak above the precursor m/z was taken as the intensity-weighted centroid above half maximum. A per-run LOESS calibration of the error against m/z was fitted in a first pass within ±40 ppm on 5,000 precursors stratified over m/z and abundance, using the monoisotopic MS1 and the fragment peaks, and peaks were then searched within ±20 ppm of the calibrated position; fragment peaks were counted when their signal-to-noise ratio was at least 5. Resolution is m/z divided by the FWHM, interpolated at half maximum, where both half-maximum crossings could be measured.

### Three-species benchmark

Mixtures of K562 (human), yeast and E. coli digests were injected at 250 ng total and acquired with ZT Scan DIA (2.8 Th) and with the Zeno SWATH DIA method of Fig. 1 (100 variable windows, 12 ms). Protein groups were assigned to a species by the UniProt entry name; groups mapping to more than one species or to none were discarded. For each of five mixture pairs (1/6, 2/5, 2/4, 3/5 and 3/4) and each acquisition method separately, the precursors quantified in all eight injections of the pair were combined into protein intensities with directLFQ, without further precursor filtering; each injection was normalized to the median log_2_ intensity of its human protein groups, and the log_2_ ratio was calculated as the difference of the mean log_2_ intensities^22^. Accuracy was assessed by an unweighted linear fit of the median log_2_ ratio of each species against the expected ratio (15 points). The spread between proteins of one species was calculated as σ_IQR_ (interquartile range/1.349) of the log_2_ ratios, σ, and expressed as (2^σ^ − 1) × 100%. Protein CVs were calculated within each mixture (for mixtures quantified in two pairs, as the mean of the two CVs) and summarized as the median per species and mixture.

### Single-cell and tissue data

All single cells were searched together. Failed cells were removed with a two-component Gaussian mixture model fitted to log_10_ precursor identifications per run, excluding runs below the equal-posterior boundary between the two components. Cell-cycle scores were computed as described previously, with protein-group intensities (log_2_ summed precursor areas) for proteins quantified in at least 50% of cells and the S-phase and G2/M gene sets of Tirosh et al.; each score is the mean of the marker proteins minus that of 50 abundance-matched control proteins per marker, and was computed only when at least three markers were quantified^45^. The pseudotime angle was taken from the centered scores and rotated so that the median G1 cell lies at zero. Rank–abundance curves use, per protein group, the median log_10_ intensity over the runs in which it was quantified; only proteins quantified in at least 50% of runs were labeled. Each dissected tissue area was searched separately. In the rank–abundance curve of the 200,000 µm² dissections, markers were labeled from a predefined list of neuronal, synaptic, glial and myelin proteins (Source Data).

### High-throughput and fractionation analyses

Protein groups per minute of elution were calculated as the mean number of protein groups per injection divided by the retention-time span containing the central 99% of identified precursors (0.5th–99.5th percentile, averaged over injections). The human digest in these analyses was HEK293F. For fractionation, 100 µg of digest was separated on a 1290 Infinity III LC system (Agilent) with an ACQUITY UPLC Peptide BEH C18 column (130 Å, 1.7 µm, 2.1 × 300 mm; Waters) at 300 µL/min and 60 °C. Buffer A was 0.1% (v/v) triethylamine in water and buffer B 0.1% (v/v) triethylamine in 90% acetonitrile, both adjusted to pH 10 with formic acid. After 3 min at 2% B, the gradient rose to 5% B at 5 min and to 38% B at 65 min, then to 100% B at 67 min, was held until 71 min and returned to 2% B until 85 min. Forty-eight 1-min fractions were collected and concatenated to 24, 12 or 6 pools by combining every 24th, 12th or 6th fraction. Human (HEK293F), yeast and E. coli digests were fractionated identically, and each pool was diluted to a nominal 250 ng per injection. Cumulative identifications of fractionated samples are the union over fractions in acquisition order.

### Differential abundance

Protein groups quantified in at least three of four samples per condition were compared on log_2_ intensities without imputation using an empirical Bayes moderated t-test following the limma model, implemented in Python (Code availability), with Benjamini–Hochberg correction^46^. Proteins with |log_2_ fold change| > 0.585 and adjusted P < 0.05 were considered to differ beyond technical variation.

### ZT Scan DIA data processing and Q1 precision

For the Q1 analyses, ZT Scan DIA files were converted to HDF5 (Hierarchical Data Format version 5) as profile data with a research-grade API for ZT Scan files; the converted files are deposited (Data availability). In the retention time × Q1 map, each cell is the summed intensity of one MS2 spectrum, with cycles assigned from the MS1 survey scans. To locate precursors in the Q1 dimension, each run was read in 20 Th Q1 blocks and divided into tiles of 11 cycles; within a tile, spectra were summed as above, centroided and the 20,000 most intense peaks kept, each with its profile across Q1. Every identified unmodified precursor was placed in the tile nearest its retention time. Theoretical b and y ions (200–1,800 m/z) that differed by at least 0.05 Th from the fragments of all other identified precursors in the same 20 Th Q1 block eluting within ±0.06 min of the tile were matched within ±0.02 Th, and the Q1 position of each fragment was taken as the vertex of a parabola through the three samples around its profile maximum. Fragments were accepted if their signal-to-noise ratio (profile maximum over the median of samples more than 1.3 Th away) was at least 5 and their cosine similarity to the precursor’s consensus profile at least 0.95; precursors with at least four accepted fragments were placed at the median of their fragment positions. The locating error is the difference between this centroid and the precursor m/z from MS1. For each isolation width, systematic trends were removed by subtracting binned medians against m/z and then retention time, and the error is reported as σ_IQR_ (interquartile range/1.349), with 95% confidence intervals from 500 bootstrap resamples. Unless stated otherwise, analyses were restricted to precursors at or above the median abundance of the run (Supplementary Fig. 2b). The Q1 axis calibration, used only for these analyses and not for the PEAKS search, was fitted with a linear retention-time term and a LOWESS curve (span 0.10) through binned medians of the deviation against Q1 position; its performance was evaluated on peptides not used for fitting (split by sequence). Transfer between injections was tested by applying an axis fitted on one injection of the 2.8 Th method to the held-out peptides of a second injection. Axes were fitted for each run. Co-isolation was counted per precursor, as the number of other precursors at or above median abundance within a window of the isolation width centered on it (or within ±1σ of the locating error) and within ±0.05 min, with precursors pooled over all acquisitions of the series; the mean and interquartile range over precursors are reported. The window-only reference was measured on the same precursors as σ_IQR_ of their distance from the center of contiguous windows of width w, over 50 window offsets. The window in Fig. 2e was analyzed in the same way, with the fragments unique among its four precursors; its chimeric MS2 spectrum is the sum of the spectra within the 2.8 Th range.

### Targeted quantification

MRM^HR^ data were read from the vendor files with alpharaw (0.4.8) without limiting the number of peaks per spectrum and processed as profile data. Scans were assigned to each target by the Q1 mass set in the method (tolerance 0.004 Th) rather than by the isolation window recorded in the file. Product-ion chromatograms were extracted at ± half the measured fragment peak width in each file (approximately ±15 ppm) and the transitions of a target were summed. The peak was located on the most abundant isotopologue (500-point interpolation, minimum signal-to-noise 3), and every level was integrated by the trapezoidal rule over ±1.5 FWHM around its own apex, located within ±1.5 FWHM of the anchor, after subtraction of an off-peak baseline (mean of the trace outside the peak region).

For the five detected peptides of the Promega 6×5 standard, transitions were fixed before quantification as the six most intense singly charged y ions of at least four residues, measured on the highest isotopologue of a reference injection and keeping only ions with at least 5% of the intensity of the most intense one; the same ion set was extracted at each isotopologue’s own product m/z. A level was accepted if the mean Pearson correlation of its summed trace with that of the highest level across the four injections was ≥ 0.80. Accepted levels had to be consecutive from the highest level down, and a peptide required at least three of them. Calibration curves were fitted by 1/x²-weighted linear least squares of peak area against nominal amount; LOD and LOQ were calculated as 3.3 and 10 times the standard error of the intercept divided by the slope. Accuracy was calculated for each injection as the back-calculated amount relative to the nominal amount, and precision as the CV across the four injections. For MS1 quantification, the first three isotopes of each isotopologue were extracted from the MS1 scans within the retention-time span of the peptide’s MS2 scans and processed identically.

For the Q1 resolution series, the calibrants at 20 fmol and above were quantified in all four runs from the y ions (1+) above the precursor m/z that differed by more than 30 ppm from the fragments of all other targets within 6 Th in Q1. Peaks were located on the highest isotopologue of each peptide and integrated as above; calibrants whose traces correlated with that of the highest isotopologue (r ≥ 0.80) in all four runs were analyzed per transition: the area of each y ion was expressed relative to the area of the same transition at open resolution. To account for co-transmitted precursor isotopes, the open-window area was corrected by the monoisotopic fraction of the complementary N-terminal part of each fragment, calculated from its elemental and isotope-label composition with alphabase.

For α-synuclein, heavy-isotope-labeled standards of the unmodified and S129-phosphorylated C-terminal peptide (JPT Peptide Technologies; C-terminal alanine labeled with ¹³C₃,¹⁵N₁) were spiked at 20 fmol per injection, and the light and heavy forms were monitored in all methionine oxidation states (12 targets). The EAD fragments z4, y4, z5 and z6 were summed for quantification; y3 was excluded for interference in its light channel; the peak was located on the heavy standard within ±8 s of the scheduled retention time and integrated as above, with the baseline taken as the 5th percentile of the extraction window where too few off-peak points remained. Areas were summed over oxidation states, and occupancy was calculated as L/H(pS129) / [L/H(pS129) + L/H(unmodified)], assuming equimolar heavy standards. For the ZT Scan DIA measurement of the same samples, the six most intense CID y ions (≥ y3, 1+ and 2+, 250–2,000 m/z) of the heavy phosphopeptide in a reference run were extracted (±25 ppm) from all spectra whose Q1 position lay within half the isolation width of each target, after excluding ions within 30 ppm of an ion of another target less than 2.5 Th away in Q1, and integrated as above. Annotated MS2 spectra were summed over the scans between the half-maximum points of the summed fragment chromatogram and matched to theoretical fragment ions within 10 ppm.

### Chromatographic peak width

Peak widths were measured on the 50% most abundant precursors of each of the four runs per method from the summed b and y fragment ions above the precursor m/z (±20 ppm, at least three fragments). Peaks were picked with an off-peak baseline and 500-point interpolation (minimum signal-to-noise 5) and accepted if the apex lay within 2 s of the retention time reported by PEAKS. Points across the FWHM were calculated as FWHM divided by the cycle time.

### Enhanced sensitivity mode comparison

Intensity ratios between acquisitions with enhanced sensitivity mode on and off were calculated from PEAKS normalized areas (mean per condition) for precursors identified in both, without cross-normalization.

## Statistics and reproducibility

Replicates are technical replicates, repeated injections of the same digest, unless stated otherwise; for single cells and tissue they are individual cells and dissections. Differential abundance between the PFF-treated and untreated cells, one culture per condition digested once and injected four times (technical replicates), was tested with a two-sided moderated t-test with Benjamini–Hochberg correction; pS129 occupancy was not tested. No data were excluded except as described above.

## Use of AI tools

Large language model–based tools (Claude, Anthropic) were used to assist with language editing of the manuscript and with drafting and debugging of the Python analysis code. All code was reviewed and tested by the authors, all reported values were verified against the underlying data, and the authors take full responsibility for the content and for the accuracy of all results. Figures 4a, 4e, 5a and 6e were made with BioRender software.

## Code availability

The analysis code, including the analysis library it bundles, is available at https://github.com/MannLabs/zenotof8600-paper; it rebuilds every figure from the deposited data.

## Acknowledgements

We thank Bioinformatics Solutions Inc. for providing access to the PEAKS software and technical support. We are also thankful to our colleagues at the Max Planck Institute of Biochemistry and SCIEX for helpful discussions and support. This study was supported by the Max Planck Society for the Advancement of Science and the Michael J. Fox Foundation for Parkinson’s Research. L. H. was supported by a Postdoctoral Fellowship of the Peter & Traudl Engelhorn Stiftung.

## Author contributions

T.H. designed and performed experiments, analyzed data, and wrote the manuscript. D.O., L.H., M.O., and L.G. performed experiments and revised the manuscript. M.B.L. developed processing scripts for targeted validation workflow. M.T. and N.E. performed single-cell proteomics experiments. C.S.S. performed the HEK293T cell culturing for the α-synuclein experiments. Y.S. performed Deep Visual Proteomics experiments and revised the manuscript. B.L., Y.L.B., N.B., M.G., A.Ca., P.R., R.N., A.P., I.B., J.C., A.Ch., G.I., K.T., T.T., B.S., J.W., and J.C.-P. contributed to instrument development and technical support and read and approved the final manuscript. M.M. conceived the study, supervised the research, wrote and revised the manuscript.

## Supplementary Information

**Supplementary Figure 1.**
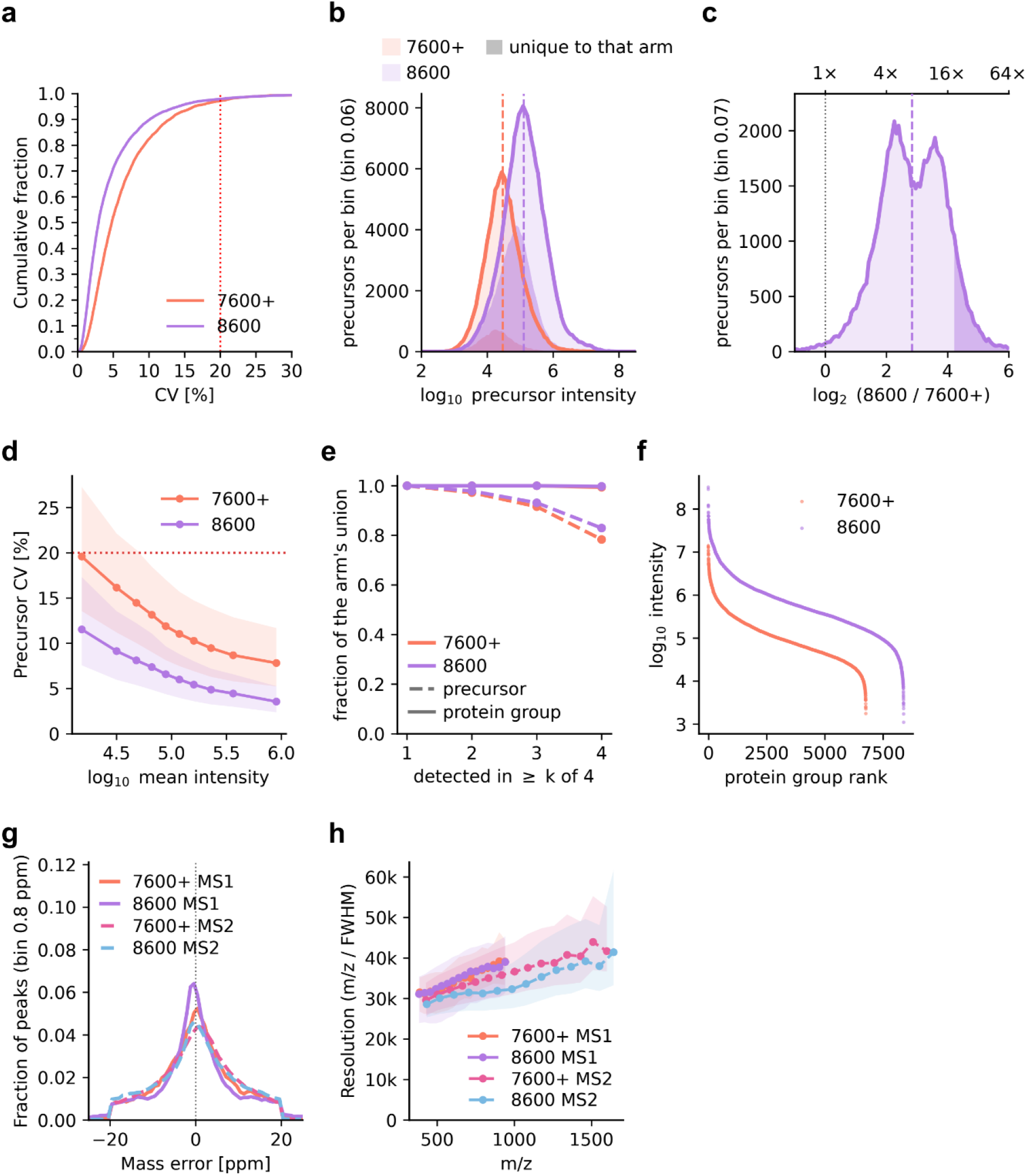
ZenoTOF 8600 system versus ZenoTOF 7600+ system, and mass accuracy and resolution. Data as in Fig. 1b–d (250 ng HEK293F digest, 60 SPD, Zeno SWATH DIA; n = 4 injections per instrument). **a** Cumulative distribution of protein group CVs; red dotted line, 20%. **b** Distribution of precursor intensity (median over injections) for all precursors identified on each instrument; darker fill, precursors identified on only one instrument; dashed lines, medians. **c** Distribution of the per-precursor intensity ratio between the two instruments for the 77,886 precursors quantified in all injections on both; dashed line, median; dotted line, no change; dark fill, top decile. **d** Precursor CV against intensity for the same precursors, in ten bins of the mean intensity of both instruments; lines, median; bands, interquartile range; red dotted line, 20%. **e** Fraction of each instrument’s identifications detected in at least k of four injections, for protein groups (solid) and precursors (dashed). **f** Rank–abundance curves of protein groups. **g** Calibrated mass error of monoisotopic MS1 precursor peaks and MS2 fragment peaks, ZenoTOF 7600+ and 8600 systems (one injection each of Fig. 1b–d; solid, MS1; dashed, MS2; Methods). **h** Resolution (m/z / full width at half maximum, FWHM) of the peaks in g whose FWHM could be measured, against m/z; lines, median; bands, interquartile range.

**Supplementary Figure 2.**
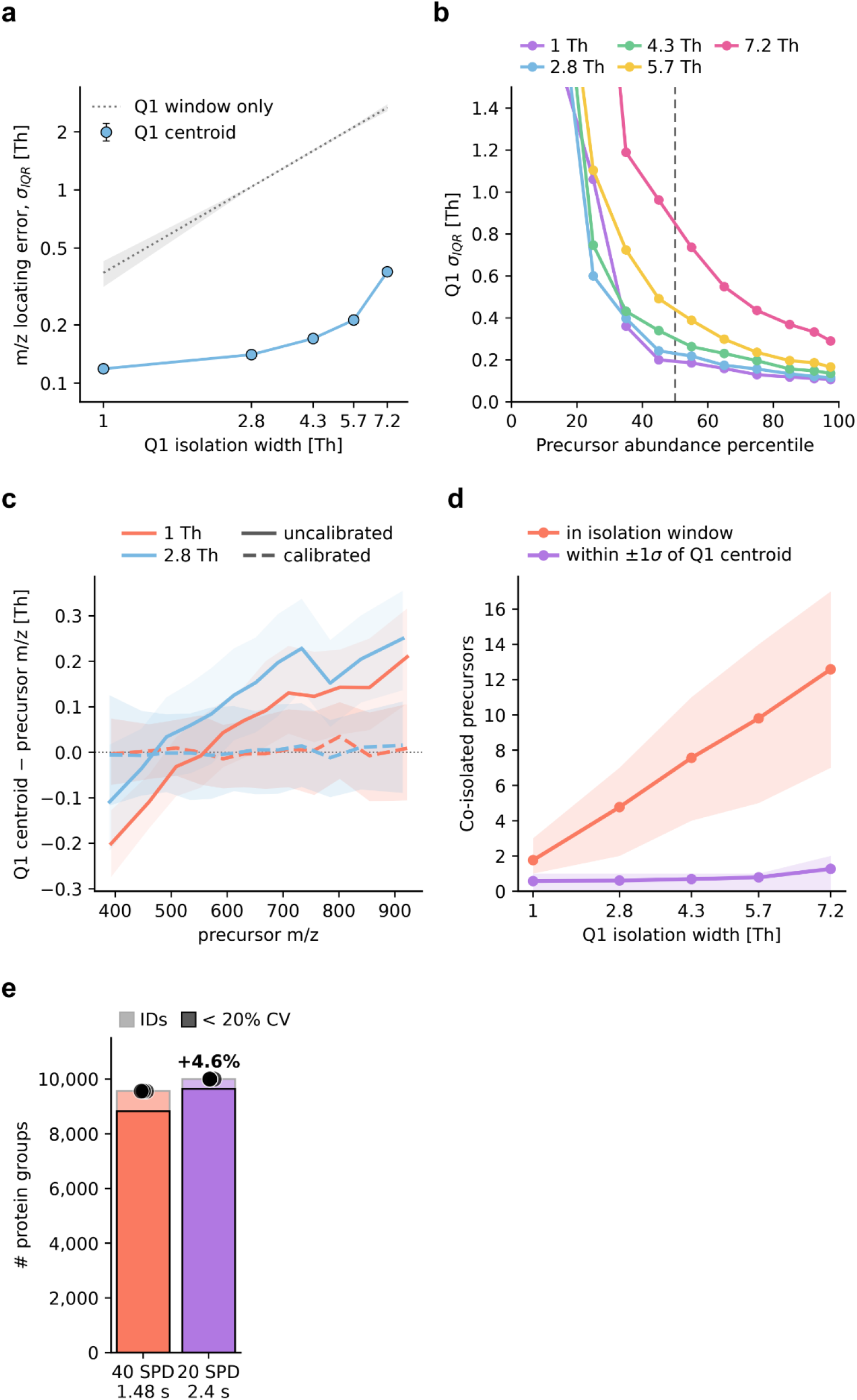
Q1 precision and acquisition speed of ZT Scan DIA. 250 ng HEK293F digest, Whisper Zoom gradient. a Locating error (σIQR, interquartile range/1.349) of the Q1 centroid against isolation width (points; error bars, bootstrap 95% confidence intervals, mostly smaller than the symbols). Dotted line and band, the error of placing each precursor at the center of its isolation window (about 0.37 × width; median and range over 50 window placements; Methods). n = 7,795–22,612 precursors per width, those with a clean Q1 profile (Methods). b Q1 locating error against precursor abundance percentile for each width. Dashed line, the median abundance used as the cutoff in a, c, d and Fig. 2; the y axis is truncated at 1.5 Th. c Systematic Q1 offset: median deviation of the precursor Q1 centroid from the MS1 precursor m/z in 12 quantile bins of m/z (about 500 and 950 precursors per bin for the 1 Th and 2.8 Th methods), before (solid) and after (dashed) Q1 axis calibration; bands, interquartile range of the individual precursors. The medians show the offset removed by the calibration; the bands show the per-precursor spread, quantified as the locating error in a. The calibration was fitted on half of the peptides and is shown for the other half. d Co-isolated precursors per precursor: other co-eluting precursors (|ΔRT| ≤ 0.05 min) within a window of the isolation width centered on each precursor (orange) and within ±1σ of its Q1 centroid (purple), against isolation width; lines, mean; bands, interquartile range. e Protein groups at 40 SPD (1.48 s cycle) and 20 SPD (2.4 s cycle) with the same 1 Th ZT Scan DIA method; n = 4 injections. Light bars, union of identifications across replicates; dark bars, subset quantified with a coefficient of variation (CV) < 20%; points, individual injections. a, c, d, precursors at or above the median abundance, placed across the whole run with identical criteria.

**Supplementary Figure 3.**
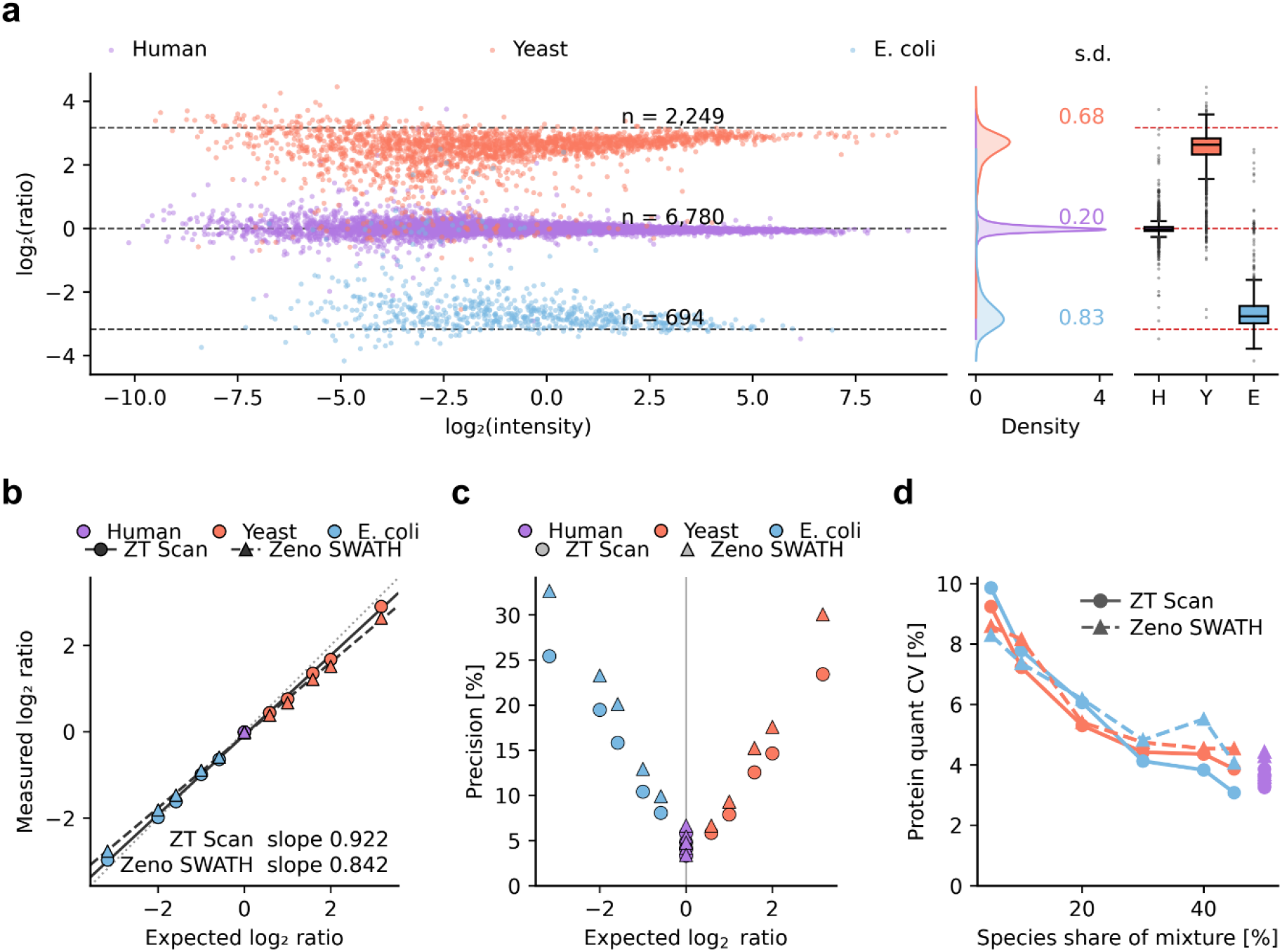
ZT Scan DIA and Zeno SWATH DIA in the three-species benchmark. Data and quantification as in Fig. 3 (n = 4 injections per mixture and method; for each pair of mixtures, proteins quantified by directLFQ from the precursors quantified in all eight injections of the pair, without precursor filtering). **a** log_2_ ratio against log_2_ intensity for mixture 1 against mixture 6, Zeno SWATH DIA, displayed as in Fig. 3e. **b** Measured against expected log_2_ ratio for ZT Scan DIA (circles, solid fit) and Zeno SWATH DIA (triangles, dashed fit); dotted line, identity. **c** Spread of the ratios between proteins of one species, (2^σ^− 1) × 100% with σ the σ_IQR_ of the log_2_ ratios, for each pair and species. **d** Median protein CV against the share of the species in the mixture.

**Supplementary Figure 4.**
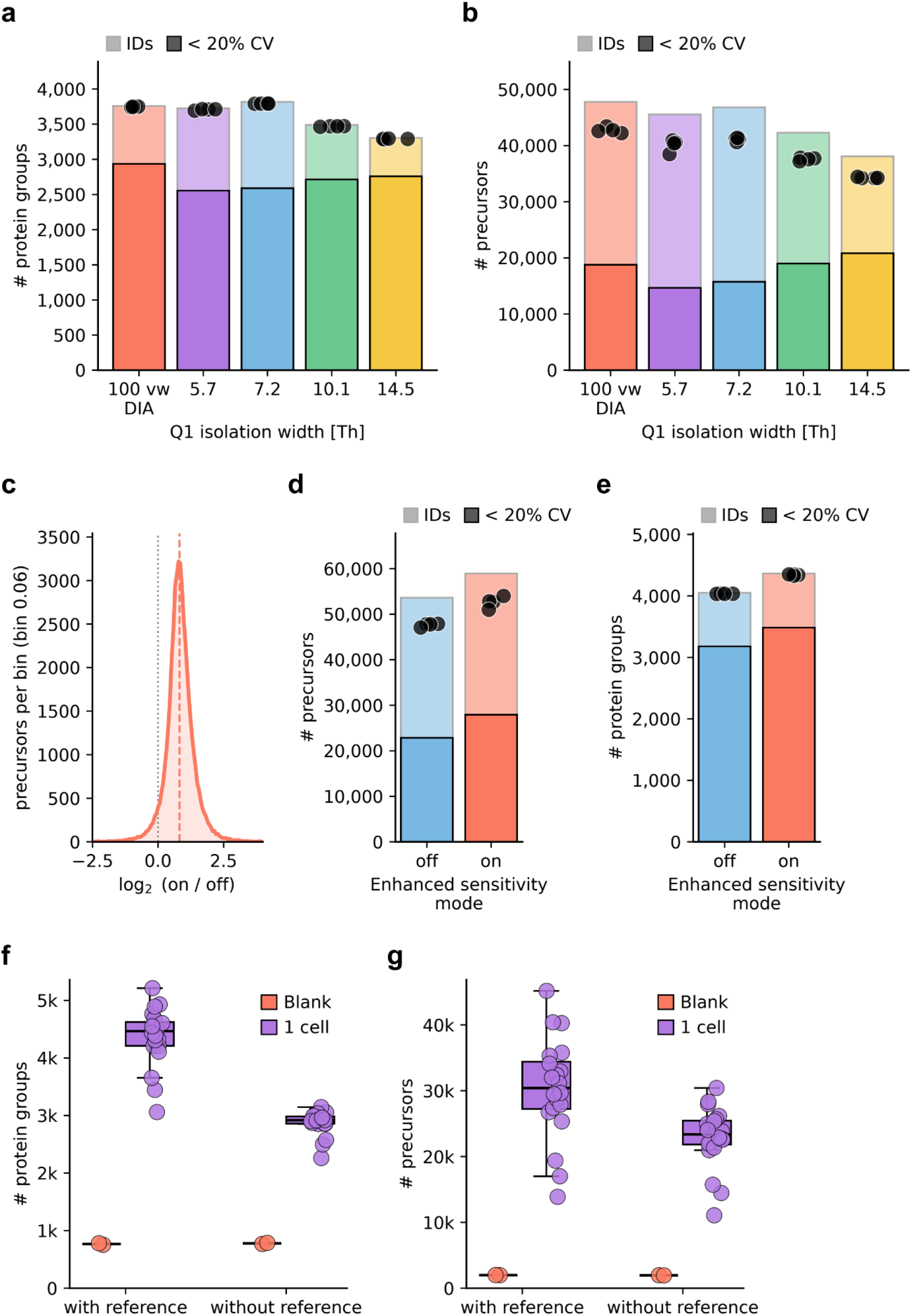
Q1 isolation width at low input, the effect of enhanced sensitivity mode and processed blanks. **a** Protein groups and **b** precursors from 0.25 ng HEK293F digest, a single-cell equivalent, acquired with Zeno SWATH DIA (100 variable windows) and ZT Scan DIA at Q1 isolation widths of 5.7–14.5 Th (1.48 s cycle, Whisper Zoom 40 SPD). Light bars, union of identifications across replicates; dark bars, subset quantified with a coefficient of variation (CV) < 20%; points, individual injections. c–e, a separate acquisition series. c Distribution of the per-precursor intensity ratio with enhanced sensitivity mode on and off (0.25 ng HEK293F digest; ZT Scan DIA, 7.2 Th, Whisper Zoom 40 SPD, the method of Fig. 4) for the precursors identified in both (n = 4 injections each); dashed line, median; dotted line, no change. **d** Precursors and **e** protein groups identified with enhanced sensitivity mode off and on (n = 4 injections each). Light bars, union of identifications; dark bars, subset with CV < 20%; points, individual injections. f Protein groups and g precursors per run for the processed blanks (n = 2) and single HeLa cells (n = 20) of Fig. 4b, searched with match between runs together with the 20-cell pools and 5 ng HeLa digest (with reference) or without them (without reference). Box, median and interquartile range of the single cells; whiskers, 1.5 × interquartile range; lines, median of the blanks; points, individual runs.

**Supplementary Figure 5.**
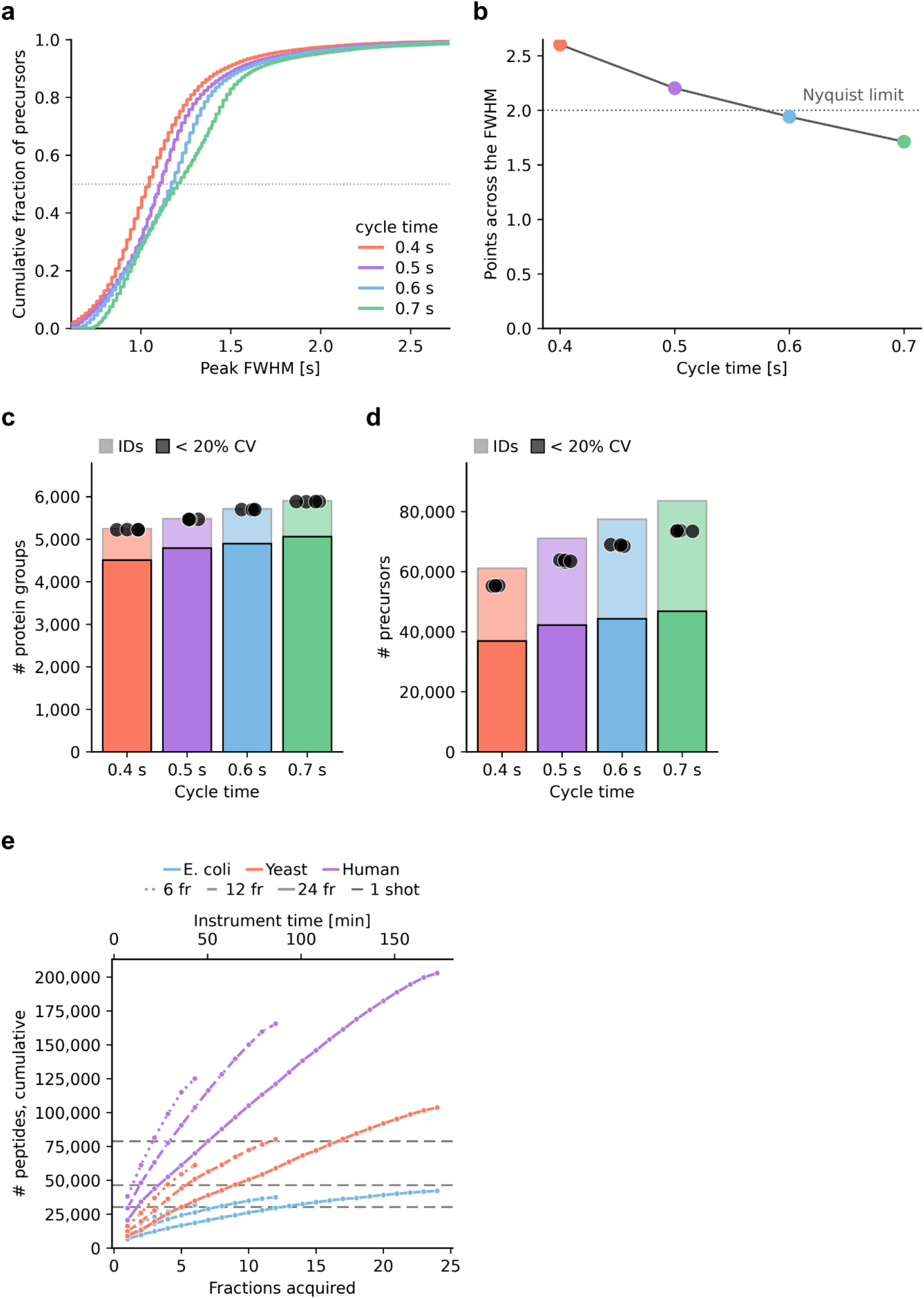
Chromatographic sampling at 500 SPD and fractionation depth. **a** Cumulative distribution of chromatographic peak width (FWHM) of the 50% most abundant precursors in each of four injections for four ZT Scan DIA methods at 500 SPD with cycle times of 0.4–0.7 s (Q1 widths 3.3–1.8 Th); dotted line, 0.5, where each curve crosses its median. **b** Median data points across the FWHM against cycle time, from the measured widths; dotted line, two points per FWHM. **c** Protein groups and **d** precursors for the four methods (n = 4 injections). Light bars, union of identifications across replicates; dark bars, subset quantified with a coefficient of variation (CV) < 20%; points, individual injections. **e** Cumulative peptides across 6, 12 and 24 high-pH reversed-phase fractions (dotted, dashed and solid lines) for human, yeast and E. coli digests; grey dashed lines, single-shot injections.

**Supplementary Figure 6.**
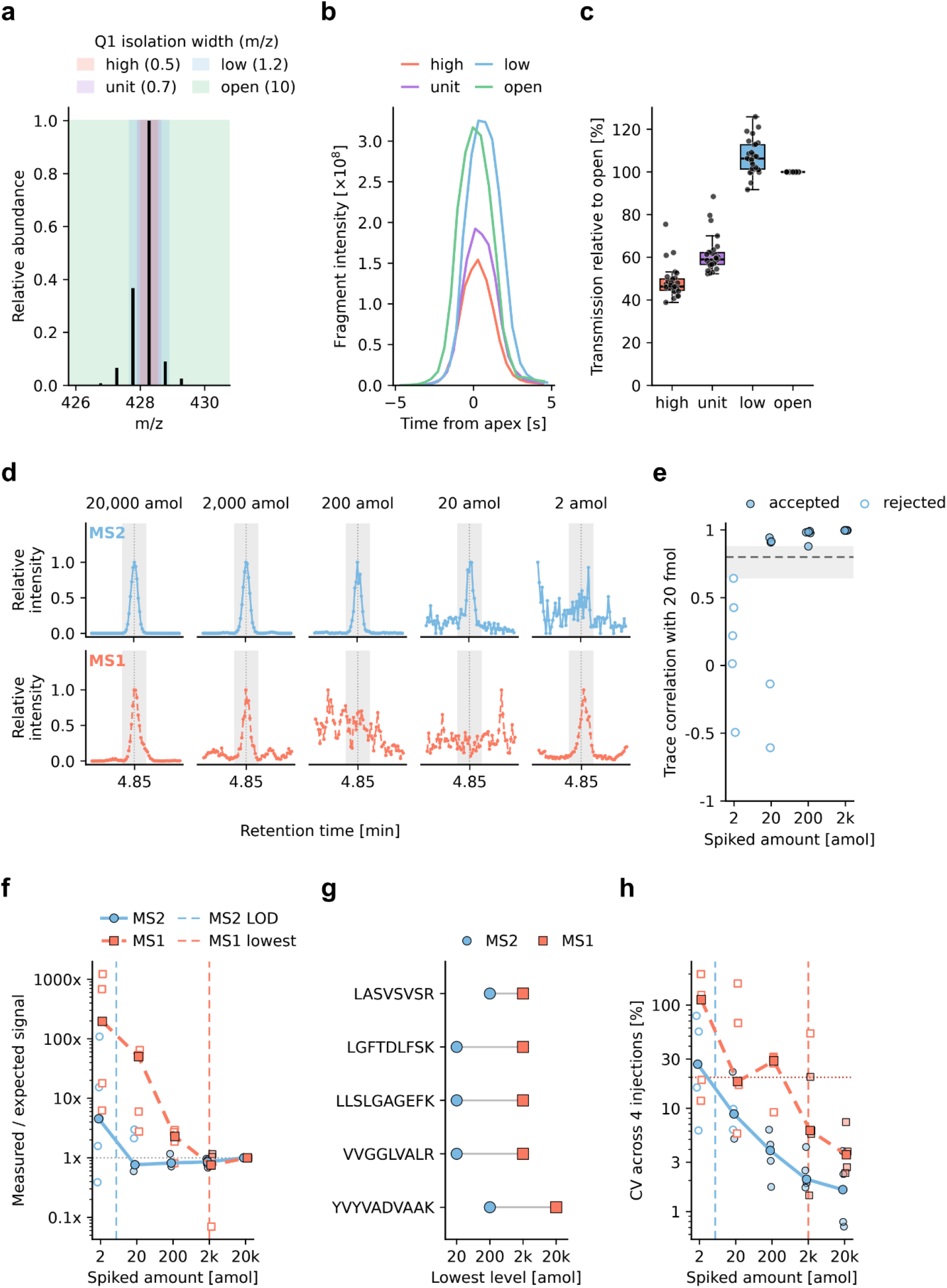
Q1 resolution and MS1 against MS2 quantification in Zeno MRM^HR^. **a** Isotope envelope of LASVSVSR (2+) against the isolation widths of the four Q1 resolution settings. **b** Summed fragment-ion chromatograms of LASVSVSR at each setting. **c** Transmission at each setting relative to open resolution, per transition (points; 29 y ions from the 9 calibrants of the Promega 6×5 standard at 20 fmol and above); boxes, median and interquartile range; whiskers, 1.5 × interquartile range. **a–c**, one Zeno MRM^HR^ injection per setting at 100 SPD. **d–h** The Zeno MRM^HR^ injections of Fig. 6a–d (n = 4), quantified from fragment ions (MS2) or precursor isotopes (MS1). **d** MS2 and MS1 chromatograms of LGFTDLFSK at each level, each scaled to its own maximum; grey, integration window; dotted line, apex of the highest level. **e** Mean correlation of the summed MS2 trace of each level with that of the highest level, per peptide; filled, accepted levels; open, rejected levels; dashed line, acceptance threshold (0.80); grey band, the range between the highest rejected and the lowest accepted value. The highest level (r = 1 by definition) is not shown. **f** Measured relative to expected signal against spiked amount; large symbols and lines, median of the five peptides; small symbols, individual peptides; open symbols, levels not accepted. **g** Lowest level accepted for each peptide. **h** CV across injections against spiked amount, displayed as in **f**; red dotted line, 20%. Dashed lines in **f**, **h**: median MS2 LOD and the lowest level accepted at MS1.

**Supplementary Figure 7.**
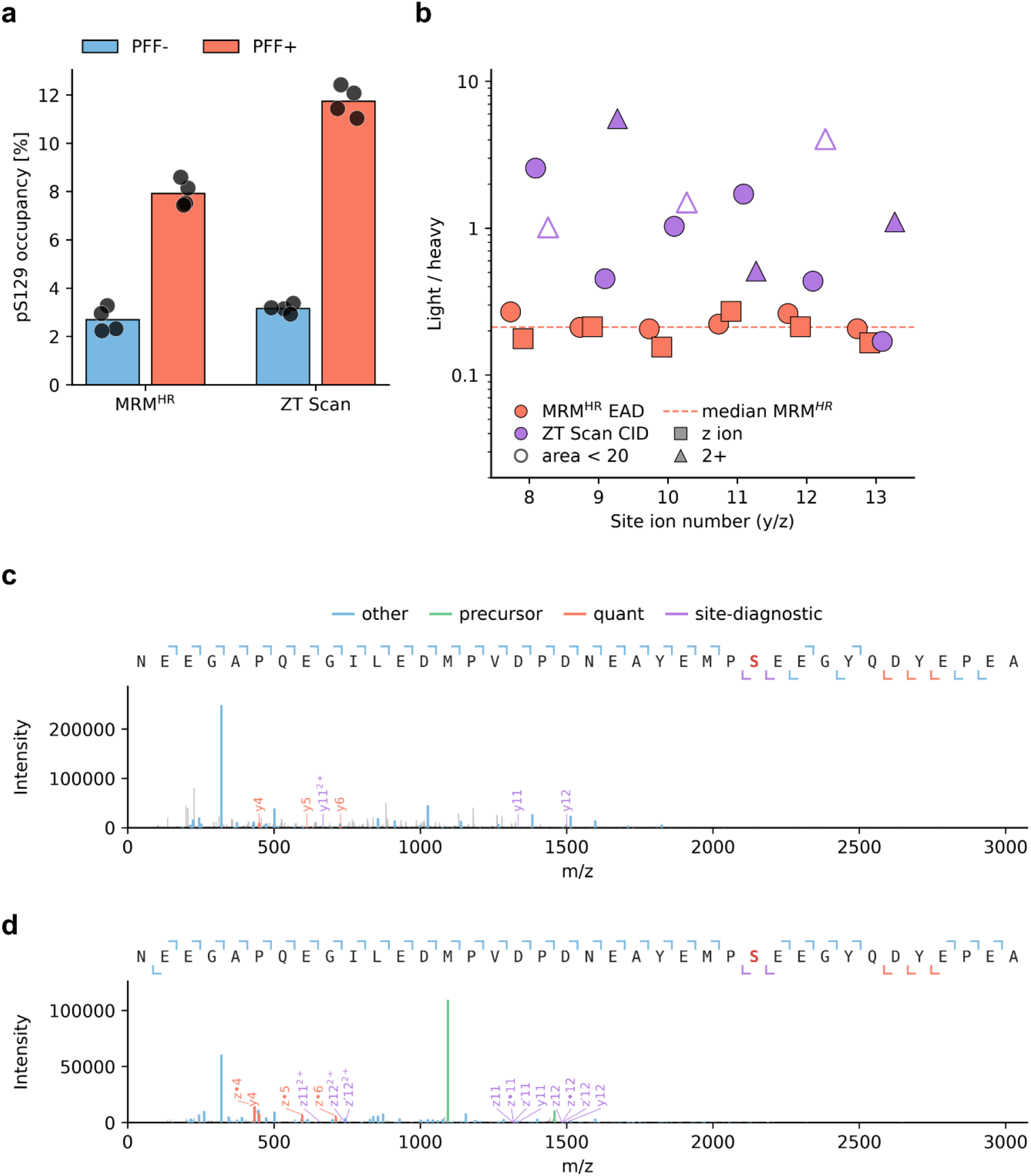
α-synuclein pS129 spectra and occupancy by two acquisition methods. a pS129 occupancy in PFF− and PFF+ HEK293T cells determined by MRM^HR^ (as in Fig. 6g) and by ZT Scan DIA from the same heavy-isotope-labeled standards; bars, mean; points, repeated injections of one digest per condition (n = 4). b Light-to-heavy ratios of the site-determining ions of the endogenous pS129 peptide in PFF+ cells, by acquisition method: EAD y8–y13 and z8–z13 by MRM^HR^ (the four PFF+ injections of Fig. 6g) and CID y8–y13 (1+ and 2+) by ZT Scan DIA (four PFF+ injections), in whose 2.1 Th window the light and heavy precursors are co-isolated; light integrated at the heavy apex and summed over injections; dashed line, median MRM^HR^ ratio; open symbols, ZT Scan DIA ions with a summed area below 20 in either channel. c, d CID (c) and EAD (d) MS2 spectra of Fig. 6h, i, with the y axes scaled to the most intense peak.

**Supplementary Figure 8.**
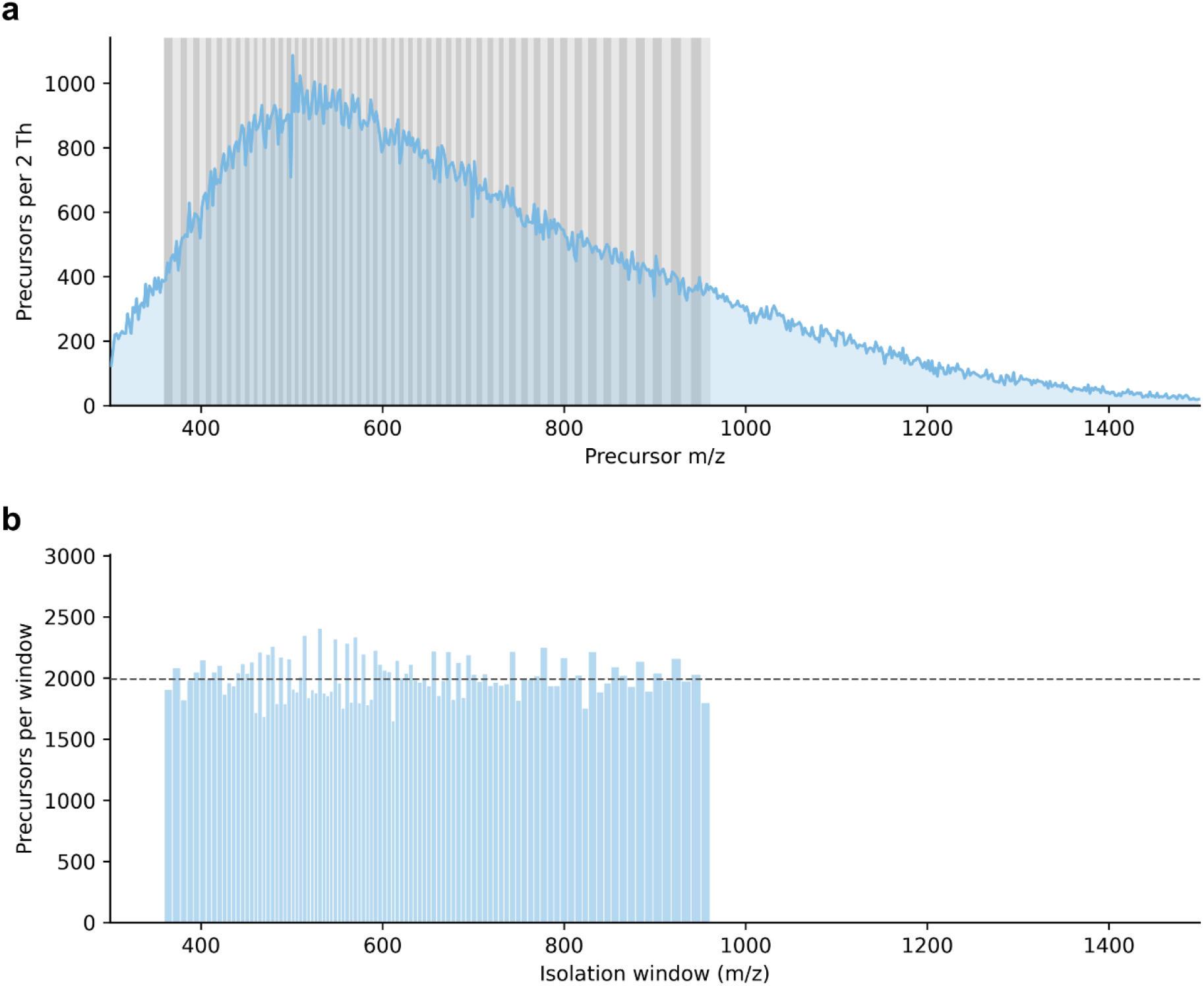
The variable-window Zeno SWATH DIA method. **a** m/z distribution of the 246,563 precursors identified by gas-phase fractionation of 250 ng HEK293F digest on the ZenoTOF 8600 system (six 40 SPD injections with 2 Th windows, each covering 200 Th between 300 and 1,500 m/z), in 2 Th bins; grey bands, the 100 isolation windows in alternating shades. **b** Precursors per isolation window; each bar spans its window; dashed line, median. The windows were designed with the dynamic binning of pyDIAid between 359 and 961 m/z and used on both the ZenoTOF 8600 and 7600+ systems (Methods).

